# Transcriptome Surveys in Silverfish Suggest a Multistep Origin of the Insect Odorant Receptor Gene Family

**DOI:** 10.1101/604389

**Authors:** Michael Thoma, Christine Missbach, Melissa D. Jordan, Ewald Grosse-Wilde, Richard D. Newcomb, Bill S. Hansson

## Abstract

The insect odorant receptor (Or) gene family is among the largest multigene families in insect genomes, but its evolutionary origin and mode of expansion is still a matter of debate. We performed transcriptomic surveys of two wingless insect species, the silverfish *Lepisma saccharina* and *Tricholepidion gertschi*, and identified multiple Or gene family members in both species. A phylogenetic analysis suggests that the silverfish Ors do not fall into the clade comprised of more derived flying insect ligand-binding Ors, but, along with bristletail, firebrat and some mayfly Ors, are consistently resolved as a distinct set of genes that may constitute an evolutionary intermediate between gustatory receptors and the more derived Ors of flying insects. We propose to consider these “primitive Ors” separately from higher insect Ors until their cellular expression patterns and function are resolved and suggest a multistep evolutionary scenario ultimately leading to the highly sensitive, rapidly evolving and physiologically diverse Or gene family observed in higher insects.

## 1. Introduction

An organism’s sensory systems are typically well tuned to the requirements of its habitat and lifestyle. Major lifestyle transitions are therefore often accompanied by dramatic changes to sensory systems, such as the secondary loss of eyes in cave-dwelling fish or the evolution of electroreception in Guiana dolphins (Cartwright et al., 2017; Czech-Damal et al., 2011; Darwin, 1859). Similarly, it is conceivable that both the conquest of land and taking flight by arthropods have created opportunities for evolutionary innovation and at the same time changed the demands on early hexapod and insect chemosensory systems.

Insects detect odorants and tastants in their environment using members of three gene families, ionotropic receptors (Irs), gustatory receptors (Grs) and odorant receptors (Ors) (Benton et al., 2009; Clyne et al., 2000, 1999; Gao and Chess, 1999; Scott et al., 2001; Vosshall et al., 1999). Ionotropic receptors are believed to have their origin early in the protostome lineage and Grs are probably even more ancient with gene family members present in animals as distantly related to insects as sea urchins (Croset et al., 2010; Eyun et al., 2017; Robertson, 2015, 2019; Saina et al., 2015). In comparison, Ors appear to be an insect-specific expansion from within the Gr gene family, with the functionally essential and highly conserved odorant receptor co-receptor (Orco) thought to be the ancestral Or (Robertson, 2019; Robertson et al., 2003).

At the molecular level, the functional Or signal transduction complex is a heteromultimer (most likely a heterotetramer) of a specific Or (Orx), which defines the ligand-specificity of the complex, and the ubiquitously expressed Orco (Butterwick et al., 2018; Dobritsa et al., 2003; Hallem et al., 2004; Hopf et al., 2015). Orco functions as a chaperone in intracellular Or trafficking and as an essential component of a non-selective cation channel that opens upon ligand-binding to the specific Orx leading to the depolarization and hence excitation of the sensory neuron expressing the Orx/Orco complex (Larsson et al., 2004; Sato et al., 2008; Wicher et al., 2008). In accordance with its central function in Or trafficking and signal transduction, exactly one Orco ortholog has been identified in most studies concerned with insect chemosensation, and insects lacking a functional Orco typically display severely impaired olfactory function (e.g. deGennaro et al., 2013; Koutroumpa et al., 2016; Krieger et al., 2003; Larsson et al., 2004; Li et al., 2016; Robertson and Wanner, 2006; Terrapon et al., 2014; Yan et al., 2017).

Two hypotheses have been proposed to explain the origin and expansion of the Or multigene family in insect genomes: (1) Ors may represent an adaptation to a terrestrial lifestyle (Brand et al., 2018; Robertson et al., 2003), which is supported by the tendency of Ors to detect hydrophobic compounds, whereas the more ancient Irs and Grs mostly detect hydrophilic substances. Moreover, Ors display, on average, a wider ligand spectrum than Irs, presumably allowing for the detection of a greater number of volatiles (Abuin et al., 2011; Ai et al., 2010; Hallem and Carlson, 2006; Min et al., 2013; Silbering et al., 2011); (2) Ors may represent an adaptation to insect flight (Missbach et al., 2014), which is supported by the observation that Ors are typically more sensitive to their ligands than Irs, that Or-expressing sensory neurons respond more readily and reliably to short odor pulses typical for in-flight odor detection than their Ir-expressing counterparts, and that sensitivity of the Or/Orco complex can be adjusted by a variety of modulatory mechanisms (Getahun et al., 2012, 2016; Guo et al., 2017; Mukunda et al., 2014; Sargsyan et al., 2011).

An apparent lack of Or gene family members in antennal transcriptomes of the bristletail *Lepismachilis y-signata* and the detection of three Orco-like sequences but no Orxs in the antennal transcriptome of the firebrat *Thermobia domestica* supported the latter hypothesis, placing the origin of the derived Orx/Orco system alongside the advent of insect flight (Missbach et al., 2014). However, this interpretation was recently challenged by the observation that one of the Orco-like sequences in the firebrat associates phylogenetically with damselfly Ors (Ioannidis et al., 2017), and even more so by the discovery of many more Or gene family members in the firebrat genome and some Or genes but no Orco ortholog in the genome of the bristletail, *Machilis hrabei* (Brand et al., 2018). Based on these findings the authors of the latter study argue that the Orx/Orco olfactory system was already fully established in the ancestor of silverfish/firebrats (i.e. the order Zygentoma) and winged insects and hence represents an adaptation to a terrestrial lifestyle.

In summary, there is growing consensus that the evolutionary origin of the Or multigene family (in the broad sense) lies in the common ancestor of insects based on transcriptomic and genomic information. However, both bristletail and Zygentoma Or repertoires display peculiarities hitherto unobserved in winged insects which appear to be specific to these wingless insect orders, with a lack of Orco in the examined bristletails and an expansion of Orco-like sequences in the firebrat. More data are clearly needed to examine whether these observations in single species generalize in the respective insect orders. Here we report results of our efforts to identify Or gene family members in the antennal transcriptomes of further examples from the Zygentoma, the common silverfish *Lepisma saccharina* and the forest silverfish *Tricholepidion gertschi*, to improve the resolution of early evolutionary events in the chemosensory systems at the base of the insect phylogeny.

## 2. Materials and Methods

### 2.1 Animals

*Lepisma saccharina* were obtained from Insect Services GmbH in September 2015 (Berlin, Germany, www.insectservices.de). *Tricholepidion gertschi* were collected in June 2016 at Angelo Coast Range Reserve, California, stored in RNALater (Ambion, Austin, Texas, USA), and sent to Jena, Germany, for further processing.

### 2.2 Dissection and RNA extraction

*Lepisma saccharina* tissues were dissected from cold-anaesthetised specimens and transferred into reaction tubes containing TRIzol reagent (Invitrogen, Carlsbad, CA, USA). Tissues were collected from ten animals each, and antennae and maxillary palps were collected from groups of mixed sex. *Tricholepidion gertschi* tissues were collected from a pool of eight individuals of mixed sex and transferred into reaction tubes containing TRIzol reagent. Tissues were homogenized using a TissueLyser LT (Qiagen, Venlo, Netherlands), and total RNA was extracted using the RNeasy Micro Kit (Qiagen, Venlo, Netherlands) and the Direct-zol RNA Micro Kit (Zymo Research, Irving, CA, USA) for *L. saccharina* and *T. gertschi* tissues, respectively.

### 2.3 Sequencing, *de novo* transcriptome assembly, quality control and quantification

RNAseq library preparation and sequencing were performed at the Max Planck-Genome-Center (Cologne, Germany). Libraries were prepared using the NEBnext Ultra Directional RNA Library Prep Kit for Illumina (New England Biolabs, Ipswich, MA, USA) and polyA enriched. For *L. saccharina*, a total number of 104,626,752 100 bp paired-end reads were generated from four different tissues (antennae: 26,609,307; maxillary palps: 26,739,878; testes: 27,056,718; ovary: 24,220,849) using the Illumina HiSeq2500 sequencing platform. For *T. gertschi*, we obtained a total of 121,012,529 150 bp paired-end reads from five different tissues (antennae: 32,823,142; maxillary palps: 32,139,161; labial palps: 19,601,531; legs: 15,056,594; body: 21,392,101). Sequencing of *T. gertschi* libraries was performed on an Illumina HiSeq3000.

For both species, an initial *de novo* transcriptome assembly was performed using the CLC Genomics Workbench 5.5 assembler (CLCbio, Copenhagen, Denmark) with default settings. In addition, we generated another assembly for each species using Trinity v2.4.0 (Grabherr et al., 2011) after adapter and quality trimming using cutadapt v1.17 (Martin, 2011) and trimmomatic v0.36 (Bolger et al., 2014), respectively. Assembly statistics for CLC and Trinity assemblies were obtained using assembly-stats v1.0.1 (https://github.com/sanger-pathogens/assembly-stats) and utility scripts contained in the trinityrnaseq package, respectively. Assembly completeness with respect to single-copy orthologs was assessed using BUSCO v3.0.2 in transcriptome mode (Simão et al., 2015) and the insect and arthropod databases. For Trinity assemblies, BUSCO analyses were performed using only the longest reported isoform per gene.

Transcript quantification was performed manually after mapping tissue-specific sequence reads to the identified and extended/curated transcripts (see below and results section) using bowtie2 v2.2.6 (Langmead and Salzberg, 2012) and with mapping results visualised in tablet v1.17.08.17 (Milne et al., 2013). Read counts (i.e. concordant mappings + single-read mappings) were normalized by the total number of reads generated per tissue to obtain fragments per million (FPM) values.

### 2.4 Or/Gr transcript identification

Transcriptome assemblies were subjected to tblastn searches using a library of insect Grs, Ors and Orcos as queries, including published wingless and Palaeoptera Ors and Grs, but also Ors from representatives of major Neoptera lineages. Contigs producing significant hits (e < 0.001) were used as queries in blastx searches against Uniprot/Swissprot to confirm them as members of the Gr/Or gene families. Identified transcripts were translated into their amino acid sequence and used as queries in iterative tblastn searches for additional Gr/Or - encoding transcripts. In addition, we screened the assemblies for candidate Orco-like sequences using hmmer v3.1 (Eddy, 2011) and a HMM profile constructed from a multiple sequence alignment of a set of insect Orco proteins.

### 2.5 Transcript extension

To obtain full length coding sequences of *L. saccharina* Or sequences, we performed Rapid-Amplification-of-cDNA-Ends (RACE) PCR using transcript-specific primers and the SMARTer RACE 5’/3’ Kit (Takara Bio USA, Inc., Mountain View, CA) according to the manufacturer’s instructions.

### 2.6 Phylogenetics

Protein sequences were aligned using MAFFT v7.271 (Katoh and Standley, 2013). After removal of misaligned sequences, the alignment was trimmed at the proposed calmodulin-binding motif in Orco (Mukunda et al., 2014) and the C-terminal fragment starting with *Drosophila melanogaster* Orco S336 was used for further analysis. Sequences containing major gaps in this region were removed from the alignment at this stage before removal of mostly empty columns using trimal v1.4.1 (Capella-Gutiérrez et al., 2009) and the -gappyout function leaving a total of 418 sequence fragments each containing 151 residues/characters. Based on the results of modeltest-ng v0.1.5 (https://github.com/ddarriba/modeltest) we used the LG model with empirically estimated amino acid frequencies and a gamma-model of rate heterogeneity with four rate categories for phylogenetic tree estimation. Maximum likelihood trees were estimated using RAxML v8.2.12 (Stamatakis, 2014), with 20 independent tree searches from random starting trees. One thousand bootstrap replicate trees were generated using the rapid bootstrap function in RAxML and transfer bootstrap support was calculated using booster v0.1.2 (Lemoine et al., 2018).

### 2.7 Data visualization

Tissue-specific expression plots were generated in R v3.2.3 (R Core Team, 2015) using the ggplot2 package v3.1.0 (Wickham, 2016). Phylogenetic trees were visualized and edited in FigTree v1.4.2 (tree.bio.ed.ac.uk/software/figtree/). All figures were edited in Inkscape v0.91 (www.inkscape.org).

## 3. Results

We performed transcriptomic surveys of several tissues in two additional zygentome species. The monophyletic order Zygentoma is the sister group to all winged insects and comprises four families (Blanke et al., 2014; Koch, 2003; Misof et al., 2014). We obtained transcriptomes from the only extant representative of the most basal family, Lepidotrichidae, the forest silverfish *Tricholepidion gertschi*, and from the common silverfish *Lepisma saccharina* from the same family as the previously examined firebrat *Thermobia domestica,* from the family Lepismatidae. Unless specified otherwise in the following sections, we will use the term Or in its broader sense (i.e. including Orco and without distinguishing between Orco and specific Or or Orx).

### 3.1 The *Lepisma saccharina* transcriptome

We generated 100 bp paired-end RNAseq data from RNA extracted from antennae, maxillary palps, ovaries and testes of a pool of cold-anaesthetised *L. saccharina* specimens. A total of ∽105 million reads were generated from the individual tissues (Table 1). These were pooled to produce two independent *de novo* assemblies using the CLC Genomics Workbench 5.5 assembler and Trinity. The CLC assembly consisted of 106,352 contigs with an N50 of 1,239 bp and an average contig size of 731 bp (Table 2). To assess completeness of the assembly with respect to gene content we performed a BUSCO analysis against the arthropod database and the results suggest that the assembly is reasonably complete and well-suited for *de novo* transcript discovery (BUSCO scores: complete: 85.7% [single: 84.1%, duplicated:1.6%], fragmented: 8.4%, missing: 5.9%). Similar BUSCO results were obtained using the insect database. The Trinity assembly consisted of 220,308 contigs with an N50 of 1,489 bp, an average contig length of 792 bp and better BUSCO scores than the CLC assembly with respect to gene content coverage (C: 97.3% [S: 96.2%, D: 1.1%]; F: 2.3%; M: 0.4%).

**Table 1.** Technical overview of sequenced transcriptomes

| Organism | Tissue | Read type | Sequencing platform | Number of reads | Sample accession |
| --- | --- | --- | --- | --- | --- |
| *Lepisma saccharina* | Antennae | 100 bp<br>paired-end | Illumina<br>HiSeq2500 | 26,609,307 | SAMN10994895 |
|  | Maxillary Palps |  |  | 26,739,878 | SAMN10994896 |
|  | Testes |  |  | 27,056,718 | SAMN10994897 |
|  | Ovaries |  |  | 24,220,849 | SAMN10994898 |
| *Tricholepidion gertschi* | Antennae | 150 bp<br>paired-end | Illumina<br>HiSeq3000 | 32,823,142 | SAMN10994899 |
|  | Maxillary Palps |  |  | 32,139,161 | SAMN10994900 |
|  | Labial Palps |  |  | 19,601,531 | SAMN10994901 |
|  | Legs |  |  | 15,056,594 | SAMN10994902 |
|  | Bodies |  |  | 21,392,101 | SAMN10994903 |

**Table 2.** Assembly summaries

| Organism | Assembler | Number of contigs | N50 | Average length | BUSCO arthropods |
| --- | --- | --- | --- | --- | --- |
| *Lepisma saccharina* | CLC | 106,352 | 1239 | 731 | C: 85.7 [S: 84.1, D: 1.6], F: 8.4, M: 5.9 |
|  | Trinity | 220,308 | 1489 | 792 | C: 97.3 [S: 96.2, D: 1.1], F: 2.3, M: 0.4 |
| *Tricholepidion gertschi* | CLC | 178,840 | 803 | 651 | C: 90.4 [S: 84.4, D: 6.0], F: 7.6, M: 2.0 |
|  | Trinity | 254,945 | 1033 | 629 | C: 93.7 [S: 83.9, D: 9.8], F: 5.1, M: 1.2 |

Using iterative blast searches, HMM searches, manual curation using the two assemblies and mapping of raw sequence reads, as well as Rapid-Amplification-of-cDNA-Ends-(RACE-) PCR with transcript-specific primers, we were able to obtain a final set of 16 transcripts encoding putative Or protein sequences ranging in length from 140 to 483 amino acids. Of these, ten are putatively full length, one is close to full length and five are partial sequences. Some of the partial Or sequences do not overlap in multiple sequence alignments and may in fact be derived from the same transcripts. Mapping of paired-end sequence reads onto the identified transcripts suggests that the partial sequences reported for Ors 12 and 13 may be derived from the same transcript representing the 5’- and 3’ ends of the Or coding sequence, respectively. We were, however, unable to verify this observation by PCR and therefore report these two sequences separately. With the exception of Or12, all the Or fragments identified in this study map to the C-terminal region of the predicted proteins and overlap in multiple sequence alignments, leading us to conclude that the *L. saccharina* genome encodes at least 15 Or gene family members.

To examine tissue-specific expression we mapped paired-end sequence reads from the tissue-specific RNAseq data onto the Or transcript set of 16 cDNAs and calculated FPM (fragments per million reads generated) values. We did not normalise to the transcript length (FPKM) in this analysis, because we assume that all Or transcripts are of similar length and normalising to the number of nucleotides would overestimate the expression of those transcripts we were unable to obtain full length sequences for. Although expression was generally low, with FPM values in the antennal transcriptome ranging from 0.04 to 8.19, all putative Or transcripts could be detected in the antennal transcriptome with at least one sequence read and, with the exception of LsacOr16, they displayed their highest levels of expression in this tissue (Fig. 1). With the exception of LsacOr15, all Or transcripts could also be detected in cDNA generated from antennal RNA by PCR using transcript-specific primers and/or by transcript-specific RACE-PCR (data not shown). In addition to their antennal expression, a subset of ten LsacOrs was also expressed in the maxillary palps. Furthermore, we observed a low level of expression of three and four Or transcripts in ovaries and testes, respectively.

**Figure 1.**
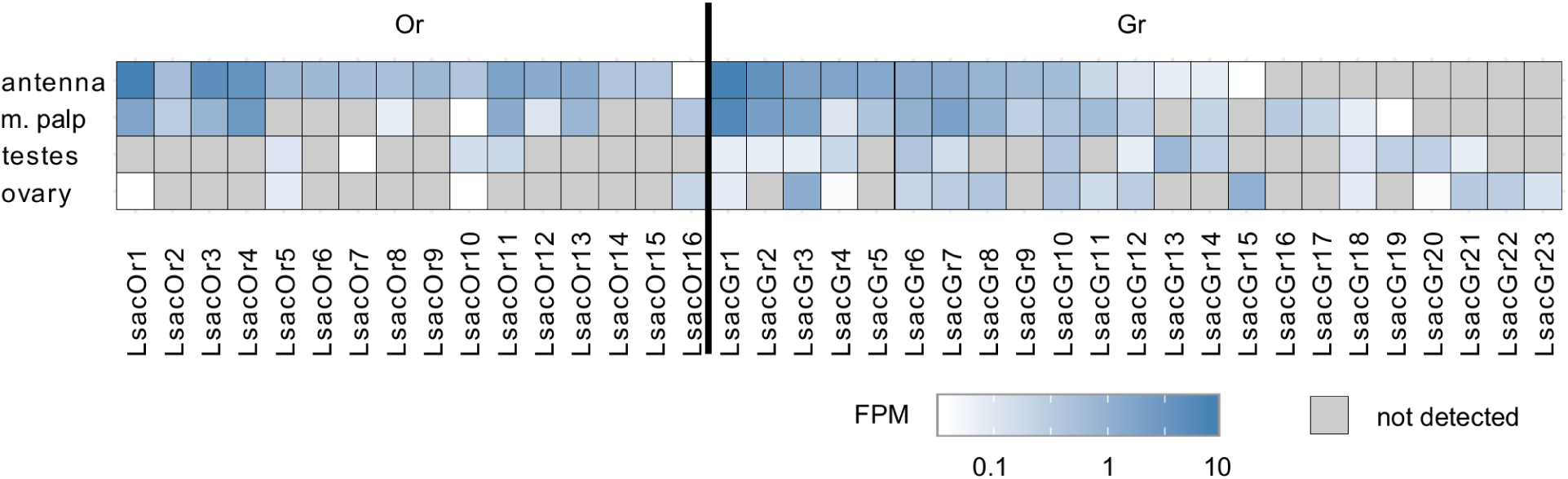
Tissue-specific expression of identified *Lepisma saccharina* chemosensory transcripts. Tissue-specific sequence reads were mapped to the final chemosensory transcripts. Expression level is presented as sequenced fragments per million reads generated (FPM).

In addition to the aforementioned Ors we were able to identify 23 candidate Gr transcripts. No attempts were made to experimentally extend these fragments by RACE-PCR, but transcript fragments were extended and curated by using the information from two independent assemblies where possible. Unlike Or transcripts, the Gr transcripts displayed greater variability in their tissue-specific expression, with some being predominantly expressed in chemosensory tissues, some in reproductive tissues and others being detected in all tissues (Fig. 1).

### 3.2 The *Tricholepidion gertschi* transcriptome

We conducted RNAseq from RNA extracted from the antennae, maxillary palps, labial palps, legs and full bodies without head of the forest silverfish *T. gertschi* (Table 1). The approximately 121 million 150 bp paired-end reads that were generated were pooled to produce two *de novo* assemblies using the CLC Genomics Workbench 5.5 assembler and Trinity. The CLC assembly consisted of 178,840 contigs with an N50 of 803 bp and an average length of 651 bp (Table 2). Trinity produced an assembly consisting of 254,945 contigs with an N50 of 1,033 bp and an average contig length of 629 bp. Despite a greater sequencing depth, the average contig lengths were lower than those obtained from *L. saccharina* in both assemblies, which hints at a higher degree of fragmentation in this dataset, probably due to a partial degradation of the RNA during storage and transport of the *T. gertschi* specimens before dissection and RNA extraction. Nevertheless, we were able to obtain high BUSCO scores for both assemblies (CLC: C: 90.4% [S: 84.4%, D: 6.0%], F: 7.6%, M: 2.0%; Trinity: C: 93.7% [S: 83.9%, D: 9.8%], F: 5.1%, M: 1.2%).

Despite the high degree of fragmentation we were able to identify 11 contigs corresponding to Or-encoding transcripts ranging in length from 269 to 1,312 bp using the same strategy employed for *L. saccharina*. Unfortunately we were unable to extend any of the candidate contigs by RACE-PCR, but some of the contigs could be improved based on the comparison of the two assemblies and mapping of raw reads. The final set of TgerOr gene family members identified in this study contains 11 partial putative Or protein sequences ranging in length from 25 to 200 amino acids. Based on sequence overlap in multiple sequence alignments we conclude that at least eight of these represent unique transcripts.

Mapping of tissue-specific sequence reads to the Or-encoding transcripts indicated that many of the contigs were assembled from single or very few fragments per tissue. Instead of providing a quantitative analysis of the tissue-specific expression levels of the identified Ors, we therefore only analysed the tissue-specific expression qualitatively (Fig. 2). As also observed in *L. saccharina*, all Or transcripts were recorded from at least one of the chemosensory tissues examined. In addition, three and five Or transcripts were also detected in the leg and body transcriptomes, respectively.

**Figure 2.**
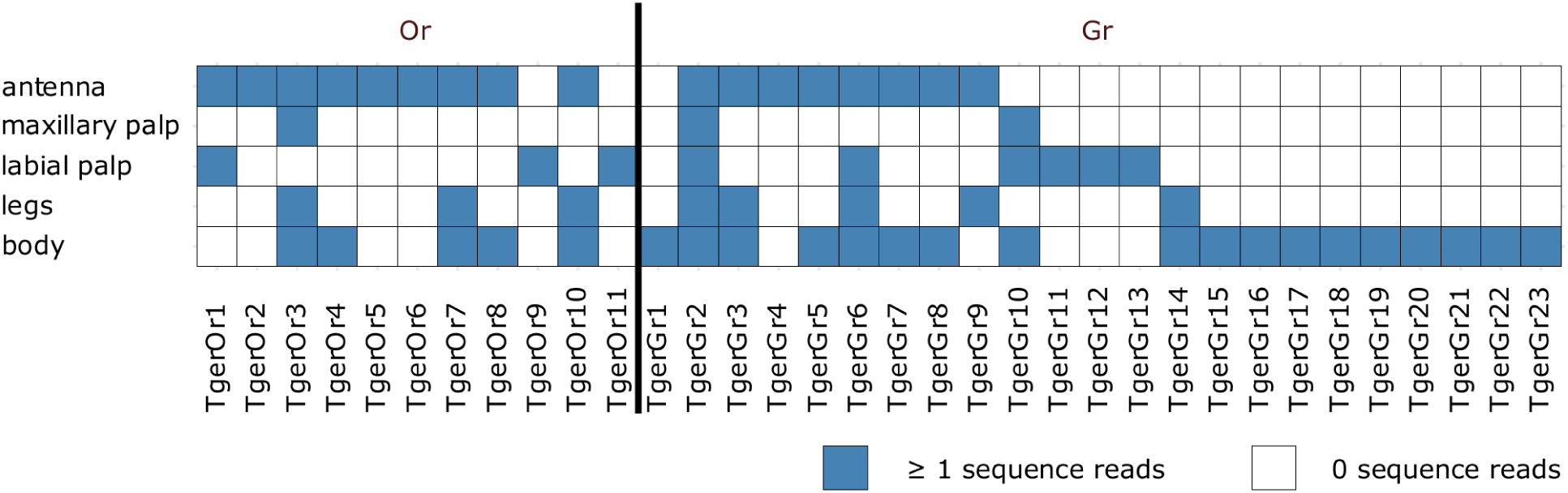
Tissue-specific expression of identified *Tricholepidion gertschi* chemosensory transcripts. Tissue-specific sequence reads were mapped to the final chemosensory transcripts and counted. Transcripts were considered present, if at least one of the tissue-specific reads mapped to the transcript.

In addition, we also detected 23 fragments of Gr transcripts, which as in *L. saccharina*, displayed greater variability in their tissue-specific expression than the Ors (Fig. 2).

### 3.3 Phylogenetic classification of Zygentoma Ors

To shed further light on early events in the evolution of the insect Or-based olfactory system and the classification of Zygentoma Ors in the context of the Gr/Or multigene family complex, we performed a phylogenetic analysis using the extracted putative chemosensory proteins and published sets of high-quality manually annotated Ors. The dataset included Zygentoma Grs used as an outgroup, bristletail Ors, Zygentoma Ors, published Palaeoptera (i.e. dragonfly, damselfly and mayfly) Ors (Brand et al., 2018; Ioannidis et al., 2017; Missbach et al., 2014) and a large selection of Neoptera (all winged insects not in Palaeoptera) Ors representing major Neoptera orders to capture neopteran Or sequence diversity. The superorder Polyneoptera was represented by the earwig *Forficula auricularia*, a member of Dermaptera, the most basally branching clade in Polyneoptera (Misof et al., 2014; Wipfler et al. 2019), for which we mined Ors from a published high-quality transcriptome assembly (Roulin et al., 2014; 1 Orco and 48 ligand-binding Ors; sequences included Supplementary Material). In addition, we included Or protein sequences from the large milkweed bug *Oncopeltus fasciatus* (Hemiptera; preprint Panfilio et al., 2017), the wheat stem sawfly *Cephus cinctus* (Hymenoptera; Robertson et al., 2018), the Colorado potato beetle *Leptinotarsa decemlineata* (Coleoptera; Schoville et al., 2018), the tobacco hornworm *Manduca sexta* (Lepidoptera; Koenig et al., 2015) and the vinegar fly *Drosophila melanogaster* (Diptera; Clyne et al., 1999; Gao and Chess, 1999; Robertson et al., 2003; Vosshall et al., 1999), as well as Orco sequences from the German cockroach *Blattella germanica* (Robertson et al., 2018a) and the fig wasp *Apocrypta bakeri* (Lu et al., 2009).

To improve our ability to assign amino acids as characters, we restricted the phylogenetic analysis to the more conserved C-terminal region of the proteins starting from the proposed calmodulin binding site, which can be aligned more confidently than the highly variable N-terminal regions (Mukunda et al., 2014; Saina et al., 2015). In addition, we employed the recently described transfer bootstrap method instead of conventional bootstrap, which is more tolerant to minor tree rearrangements and single-taxon differences than the essentially binary standard bootstrap and therefore tends to perform better than conventional bootstrap for large datasets and particularly for deep branching patterns (Lemoine et al., 2018).

The inferred phylogeny supports the monophyly of the Or gene family with respect to the Gr family with a high degree of confidence (Fig. 3). In addition, we find a highly supported monophyletic clade formed by the neopteran Ors, which does not include any of the palaeopteran or apterygote (i.e. bristletail and Zygentoma) Ors. We will therefore use the term “primitive Ors” for those protein sequences which were resolved as members of the Or multigene family, but that do not belong to the clade comprised of neopteran ligand-binding Ors (Orx). Whereas phylogenetic information on insect orders is lost within the neopteran Or lineage, primitive Ors are consistently resolved outside of and more closely related to Grs than the neopteran Ors. With the exception of a single *O. fasciatus* Or (OfasOr79), which resolves more closely to Grs than any of the other Ors, a clear separation between primitive Ors and neopteran Ors was also observed in a phylogenetic analysis using full length sequences of the proteins (Fig. S1).

**Figure 3.**
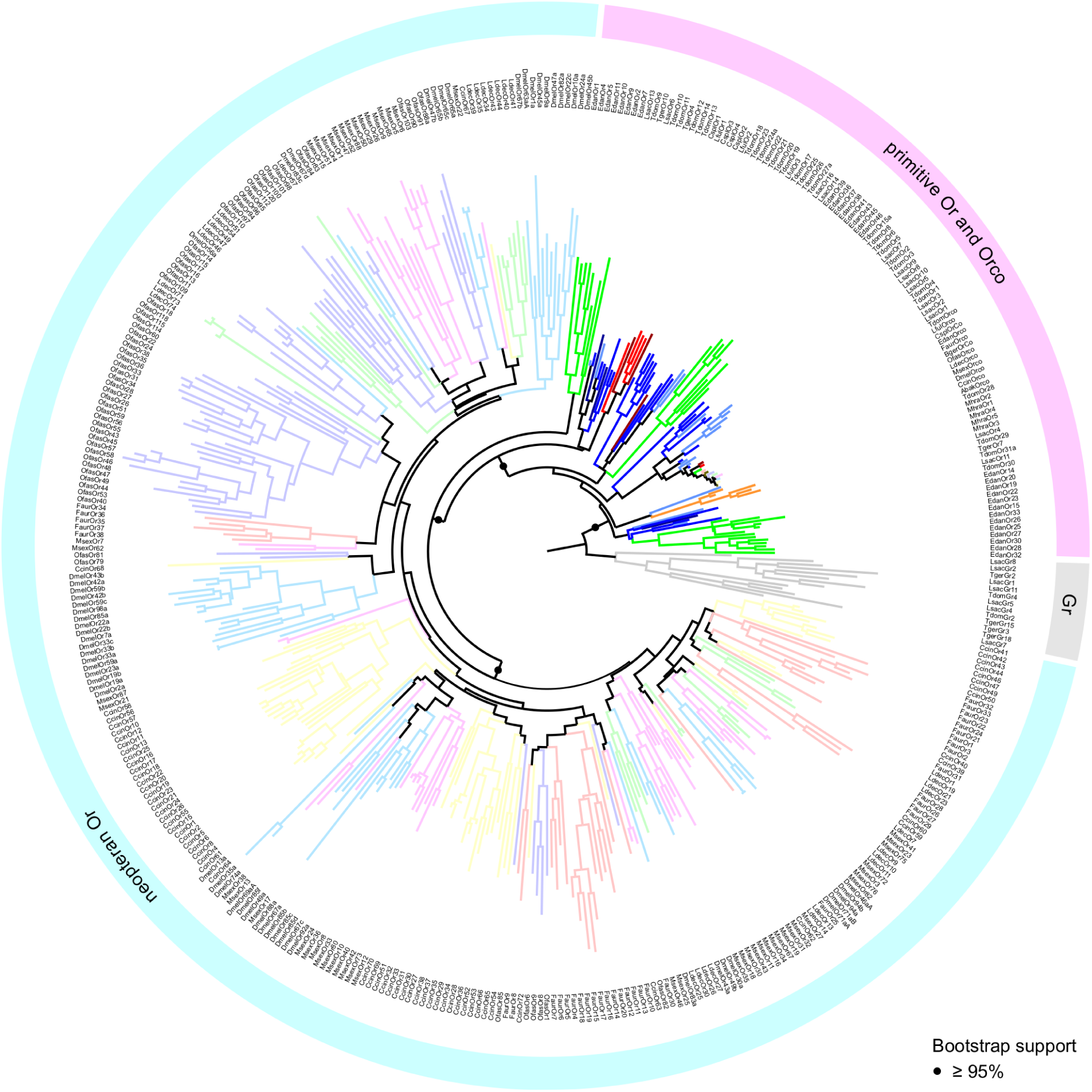
Maximum Likelihood tree of insect Grs and Ors. Tree is rooted using Grs (grey) as the outgroup. Different colours represent different species. Neopteran Ors represented by desaturated colors. Statistical support of selected nodes is presented as transfer bootstrap percentages. Species abbreviations: Lsac – *Lepisma saccharina (*light blue); Tger – *Tricholepidion gertschi* (dark blue); Tdom – *Thermobia domestica (*blue); Mhra – *Machilis hrabei* (orange); Edan – *Ephemera danica* (green); Lful – *Ladona fulva* (red); Cspl – *Calopteryx splendens* (dark red); Faur – *Forficula auricularia* (desaturated red); Ofas – *Oncopeltus fasciatus* (desaturated purple); Ccin – *Cephus cinctus* (desaturated yellow); Ldec – *Leptinotarsa decemlineata* (desaturated green); Msex – *Manduca sexta* (desaturated pink); Dmel – *Drosophila melanogaster* (desaturated blue); Bger – *Blattella germanica* (brown); Abak – *Apocrypta bakeri* (brown).

Within the group of primitive Ors we were able to recover all previously reported major clades (Brand et al. 2018). However, because we rooted the tree using Grs rather than the Orco clade as the outgroup, and included neopteran Ors to analyse palaeopteran and apterygote Ors in the broader context of the evolution of the entire Or multigene family, these clades are arranged differently in our phylogenetic construction. Each of the palaeopteran orders and the order Zygentoma contain one lineage-specific expansion which closely associates with neopteran Ors (Fig. 4). With the exception of LfulOr3 from the dragonfly *Ladona fulva*, these expansions include a highly supported clade of all dragonfly/damselfly Ors, and a mayfly-specific clade including EdanOrs 1-11 from *Ephemera danica*. The Zygentoma Or clade which closely associates with neopteran Ors comprises *T. domestica* Ors 9-14, *L. saccharina* Ors 6 and 13 and two of the *T. gertschi* Ors, TgerOr4 and TgerOr10. In this phylogenetic construction, the Orco clade does not appear to form the base of the Or gene family, but, rather, constitutes an expansion within the primitive Ors and includes all palaeopteran and neopteran Orco proteins, TdomOrco and two of the *L. saccharina* Ors, LsacOr1 and LsacOr2, but none of the *T. gertschi* Ors. This Orco clade is a member of a larger well-supported clade that includes TdomOrs 1-6 and 8 and LsacOrs 3, 5, and 7-10. The other bristletail, Zygentoma and Palaeoptera Ors form lineage- and species-specific expansions within the group of primitive Ors, but the exact sequence of evolutionary events within the primitive Ors could not be confidently determined given the current amount of sequence information. The major patterns described above with a clear separation between primitive Ors and neopteran Ors, however, remained consistent over several phylogenetic analyses using different sets of Grs and Gr-like proteins and neopteran Ors (analyses not shown).

**Figure 4.**
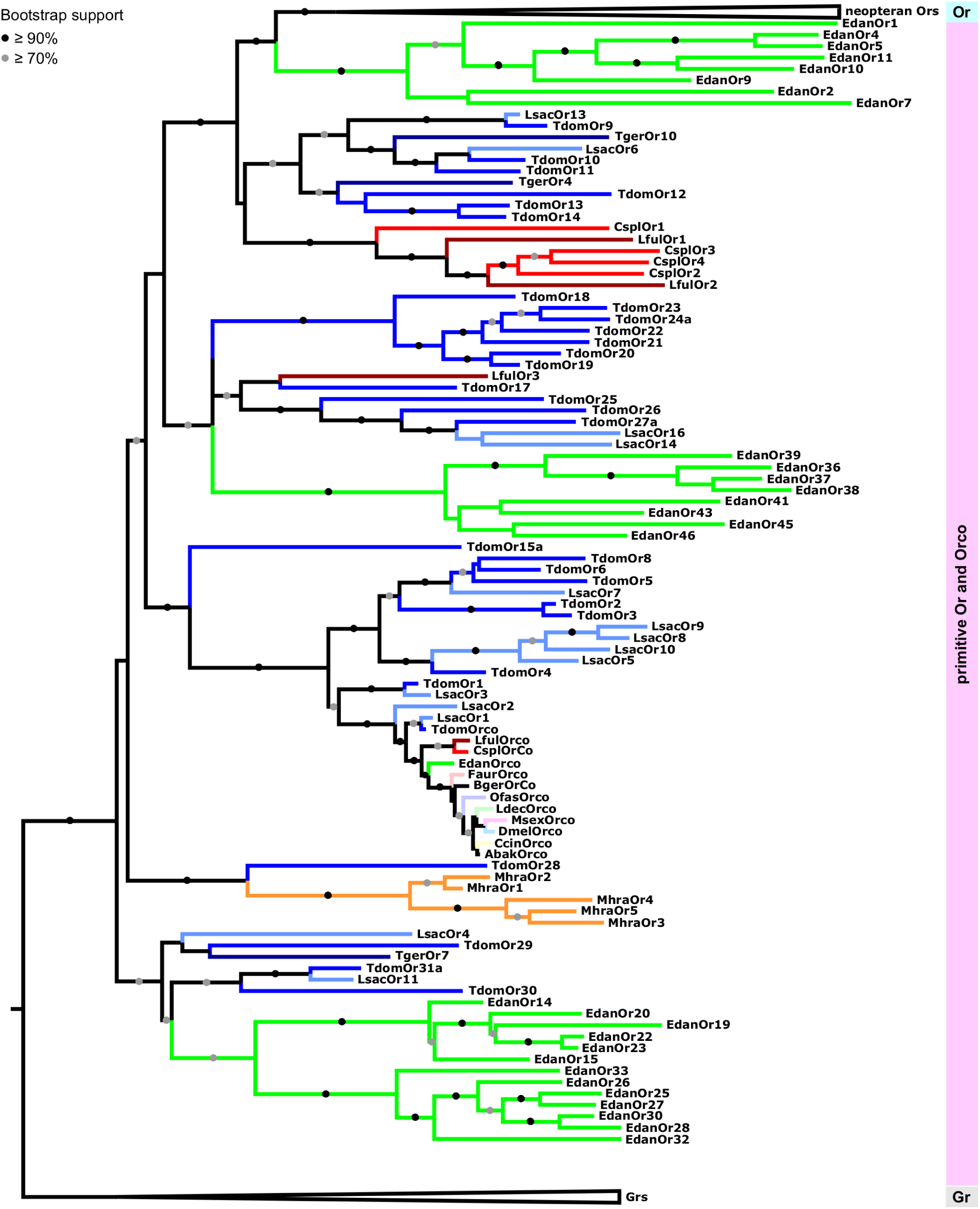
Maximum likelihood tree of Ors focussing on the primitive Or group including Orco within the phylogeny. Tree is rooted using Grs as the outgroup. Node support is presented as transfer bootstrap percentages. Different colours represent different species. Different orders are represented by different color hues. Species abbreviations: Lsac – *Lepisma saccharina (*light blue); Tger – *Tricholepidion gertschi* (dark blue); Tdom – *Thermobia domestica (*blue); Mhra – *Machilis hrabei* (orange); Edan – *Ephemera danica* (green); Lful – *Ladona fulva* (red); Cspl – *Calopteryx splendens* (dark red); Faur – *Forficula auricularia* (desaturated red); Ofas – *Oncopeltus fasciatus* (desaturated purple); Ccin – *Cephus cinctus* (desaturated yellow); Ldec – *Leptinotarsa decemlineata* (desaturated green); Msex – *Manduca sexta* (desaturated pink); Dmel – *Drosophila melanogaster* (desaturated blue); Bger – *Blattella germanica* (brown); Abak – *Apocrypta bakeri* (brown).

## 4. Discussion

Despite occupying a key position in the insect phylogeny of being the most derived wingless insect order, members of Zygentoma have historically received little attention regarding their chemosensory systems and chemical ecology. Firebrats and silverfish are known to possess a variety of cuticular sensory hairs, sensilla, on their appendages, some of which have been shown to have a chemosensory function by physiological methods (Adel, 1984; Berg and Schmidt, 1997; Hädicke et al., 2016; Hansen-Delkeskamp, 2001; Missbach et al., 2014). Behaviorally, firebrats and silverfish tend to aggregate in response to some chemical stimuli and to avoid others (Wang et al., 2006; Woodbury and Gries, 2013, 2008, 2007). In contrast, little has been known concerning the molecular makeup of their Or-based chemosensory system until a publication reported three Orco-like transcripts expressed in the antenna of the firebrat, *Thermobia domestica* (Missbach et al., 2014). The presence of three co-receptors without a single Orx seemed strange at the time, but a recent study examining the genome of the same species identified 44 members of the Or multigene family and proposed that Zygentoma already possess a full Orco/Orx-based olfactory system (Brand et al., 2018). The discrepancy between the number of Or transcripts identified from chemosensory tissue and the number of Or genes present in the genome may in part be attributable to an insufficient depth of the RNAseq experiments or point to an expression profile that involves other tissues or other life stages.

The present study provides several lines of evidence that Ors have a chemosensory, if not necessarily olfactory, function in the order Zygentoma. We find eight and 15 Or transcripts in transcriptomes from the silverfish *T. gertschi* and *L. saccharina*, respectively, numbers which are higher than those previously reported from the firebrat antenna. The 15 Ors from *L. saccharina* and eight from *T. gertschi* that we report here are likely to be underestimates of the full repertoire of these genes found within the respective genomes given that we are sampling just a small number of tissues and that there are often low expression levels of these genes per tissue due to their expression being limited to certain sensory neuron types. Consistent with an olfactory function and unlike identified Grs, the Or transcripts were predominantly detected in chemosensory tissues such as antennae and maxillary palps, and some of them were also expressed in other tissues. Expression of Ors outside the classical chemosensory tissues has also been reported in other insect species (Bray and Amrein, 2003; Haverkamp et al., 2016; Klinner et al., 2016; Pitts et al., 2014; Raad et al., 2016). Given the generally low expression levels of Ors and that many of the firebrat Ors identified from the genome form clades with silverfish Ors identified from our RNAseq experiments in our phylogenetic analysis, it is likely that many of the firebrat genes described from the genome by Brand and co-workers (2018) are indeed also expressed in the firebrat antennae and that they were missed in previous RNAseq experiments because of insufficient sequencing depth.

Both insect Ors and the phylogenetically unrelated vertebrate Ors are assumed to evolve rapidly following a birth-and-death evolutionary process with new genes being born by gene duplication and others being lost by pseudogenisation (Andersson et al., 2015; Benton, 2015; Hansson and Stensmyr, 2011; Hughes et al., 2018; Nei et al., 2008; Nei and Rooney, 2005; Ramdya and Benton, 2010; Sánchez-Gracia et al., 2009). In combination with the adaptation of the Or repertoire to the species’ ecological niche, this evolutionary process leads to Or gene phylogenies mostly lacking clear orthologs between species, and with species-specific expansions and reductions, which do not necessarily follow the underlying species phylogeny. Such a branching pattern with species-specific expansions and a lack of clear orthologs between species is clearly apparent in our phylogenetic reconstruction of neopteran ligand-binding Ors. Although the numbers of wingless and palaeopteran species, for which Or gene/transcript sequences are available, is still limited, a similar pattern begins to emerge within the group of Zygentoma Ors. This pattern on the one hand further supports the hypothesis that primitive Ors may function as true odorant receptors in these insect species, as they appear to follow an evolutionary process that is indicative of rapid evolution in response to the demands of the organism’s ecological niche. On the other hand it suggests that Zygentoma may have a much richer species-specific chemosensory ecology than they have historically been given credit for.

But what does the Zygentoma Or signal transduction complex look like? Based on the finding that the firebrat genome harbours one Orco-like sequence that is more similar to neopteran Orcos than the others, the authors of a previous study concluded that the firebrat possesses an olfactory system akin to that of more derived insects consisting of one universal co-receptor and multiple ligand-binding Ors (Brand et al., 2018). Although such a molecular makeup of the zygentome olfactory system may indeed be true, some of our findings as well as results from a previous study suggest that the expression logic and molecular function of the zygentome Or signal transduction complex is likely to differ from that of the Neoptera (Missbach et al., 2014). Firstly, in the firebrat, there is a stark contrast in the number of olfactory sensory units predicted from the Or gene number, when assuming a conventional Orx/Orco system with one universal co-receptor, i.e. 43 olfactory sensory neuron types, and those observed in physiological experiments, i.e. 13 functional units. Of those 13 functional units in the firebrat antenna, six display tuning profiles typically observed from Ir-expressing sensory neurons, leaving seven putative Or-expressing sensory neurons to house 43 possible Orx/Orco combinations predicted from genomic information. While it is possible that some of the firebrat Ors are expressed in different tissues or at different life stages, or that the electrophysiological screen simply missed some sensilla or sensory neuron types, this difference in numbers is striking. In our phylogenetic reconstruction we observed an expanded Orco clade which contains all winged insect Orco proteins, as well as eight firebrat Ors (including the proposed sole Orco) and eight *L. saccharina* Ors. The firebrat Ors in this clade share an intron-exon-structure with the neopteran Orco (Brand et al., 2018), and all of the Zygentoma Ors in this expanded Orco clade display a higher sequence identity with neopteran Orco proteins than that typically observed for neopteran ligand-binding Ors. This higher level of sequence conservation may indicate some degree of functional conservation. We therefore propose that the Zygentoma olfactory system does not possess one universal, but rather several co-receptor proteins. The functional Or signal transduction complex in firebrats and silverfish will then consist of stereotypic combinations of co-expressed Ors with one of the members of the expanded Orco clade being one of the complex constituents. Such an expression logic would be similar to that of the Gr gene family from within which Ors are thought to have originated (Dahanukar et al., 2007; Jiao et al., 2008; Jones et al., 2007; Kwon et al., 2007; Lee et al., 2009; Moon et al., 2009).

Including both Grs and neopteran Ors in our phylogenetic analysis allowed us to examine branching patterns within the insect Or multigene family at greater detail than has been possible previously. Although both primitive and neopteran Ors appear to evolve rapidly and presumably follow a birth-and-death evolutionary process, they were consistently resolved as two separate non-overlapping groups of genes in phylogenetic reconstructions including different sets of Grs and neopteran Ors. Our analyses thus suggest, that the Or multigene family found in neopteran insects is the result of not one, but two rapid expansions. The first ‘primitive Or’ expansion presumably occurred in the common ancestor of insects in the mid-to-late-Ordovician (Misof et al., 2014; Rota-Stabelli et al., 2013). At this time the first vascular land plants had already emerged (Steemans et al., 2009) adding complexity and novel compound classes to the chemical landscape surrounding early terrestrial arthropods. Given the unclear positioning of the Orco clade in our phylogenetic reconstruction and because some Grs are able to form functional homomultimeric complexes in heterologous systems (Sato et al., 2011; Zhang et al., 2011), it is not clear at this point whether the Or gene family originated from a set of Gr genes, whose gene products functioned as heteromultimers, or from a single Gr gene. However, the second alternative seems attractive, because it can easily accommodate the as yet enigmatic absence of an Orco ortholog from the bristletail genome.

The co-receptor Orco most likely originated in the common ancestor of Zygentoma and winged insects (Brand et al., 2018). Given that just one Orco has been described from all winged insects (i.e. Palaeoptera and Neoptera) so far, the expansion of Orco-like sequences potentially represents a derived character of the Zygentoma lineage and not the ancestral state of the insect olfactory system. The absence of Orco in bristletails, however, suggests that some primitive Ors may have functioned and may still be functioning without a co-receptor or in variable heteromultimers not only in bristletails, but also in Zygentoma and Palaeoptera.

Primitive Ors are mostly absent in damselflies and dragonflies apart from the clade closely associating with neopteran Ors, which may well constitute the clade of ligand-binding Ors associating with these species’ Orco proteins. However, they are still present with large expansions in the mayfly *E. danica*. According to our phylogenetic reconstruction, primitive Ors were entirely lost from insect genomes at the base of Neoptera in the early Devonian (Misof et al., 2014; Rota-Stabelli et al., 2013), leaving only Orco and one or a few Orx genes. The gradual loss of all Or genes except Orco and the Orxs forming complexes with Orco (i.e. the ancestral ligand-binding Ors) thus correlates with the advent of insect flight and the raised demands of in-flight odour detection. The gene products of *T. domestica* Orco and *L. saccharina* Or1 already possess some of the amino acid residues and motifs identified as targets of the molecular mechanisms modulating the sensitivity of the Orco/Or complex such as potential phosphorylation sites and a proposed calmodulin binding motif (Mukunda et al., 2014; Sargsyan et al., 2011). It is therefore tenable that the complex of Orco and the ancestral Orx replaced primitive Ors at the advent of flight, because it enabled early flying insects to detect the faintest whiff of an odour possibly indicating a valuable resource while still being able to adapt to the higher odor concentrations present at the odor source. After the loss of primitive Ors the radiation of neopteran insects in the Devonian then led to a second expansion of the Or multigene family giving rise to the large family of Ors observed in higher insects today.

In conclusion, we propose that the origin of the derived Orx/Orco-based insect olfactory system lies in a sequence of: (1) an initial rapid diversification of primitive Ors starting in the common ancestor of insects, which gave rise to the universal co-receptor and the ancestral ligand-binding Or/s at some later point; (2) pruning of this initial Or expansion was mediated by the demands of in-flight odor detection; (3) another rapid expansion and diversification of the remaining Orx lineage in Neoptera. Thus, both terrestrialisation and the advent of flight likely had dramatic effects on the molecular makeup of hexapod chemosensory systems. The proposed scenario will need to be put to the test by further increasing the phylogenetic resolution at the origin of the Or gene family with the addition of more genetic information on Ors from wingless insects and Palaeoptera and, importantly, Grs from non-insect hexapods, which may close the gap between the Gr and Or gene families. Aside from adding further sequence information, future studies should be directed at elucidating the expression logic and function of primitive Ors. The chemosensory systems of bristletails, silverfish and mayflies may well afford us with the rare opportunity to catch a glimpse of the evolutionary past and to understand the intermediate steps in the evolution of a multigene family not only from the sequence perspective, but also at a functional level.

## Funding

This study was supported by the Max Planck Society, a Royal Society Marsden Fund grant to RDN and MDJ (15-PAF-007) and a DFG Research Fellowship awarded to MT (TH 2167/1-1).

### Acknowledgements

We thank Dr Markus Koch, Prof. Mary Power, Prof. Kipling Will and Peter Steel for support in finding a source for *T. gertschi* specimens, Dr Joyce Gross and Diane M. Erwin for collecting the animals, and the Angelo-Steel family and the University of California Natural Reserve System for providing a protected site for the research. We are grateful to Dr Jean-Claude Walser for granting us access to the *F. auricularia* transcriptome. We thank Sascha Bucks for support with RNA extractions and Domenica Schnabelrauch for sequencing of PCR products. We also thank Sridevi Bhamidipati and Dr Colm Carraher for comments on the manuscript.

## Author contributions

This study was conceived by all authors. Data collection and analysis was performed by MT, CM and EGW. Writing of the initial draft of the manuscript was performed by MT. All authors contributed to revising the manuscript and the acquisition of funding.

## Conflict of interest

The authors declare that the research was conducted in the absence of any commercial or financial relationships that could be construed as a potential conflict of interest.

## Data availability

Raw sequence reads have been submitted to the NCBI short read archive as BioProject PRJNA523997. Transcriptome assemblies will be provided upon request.

