## Supplementary Material for "Transcriptome Surveys in Silverfish Suggest a Multistep Origin of the Insect Odorant Receptor Gene Family"

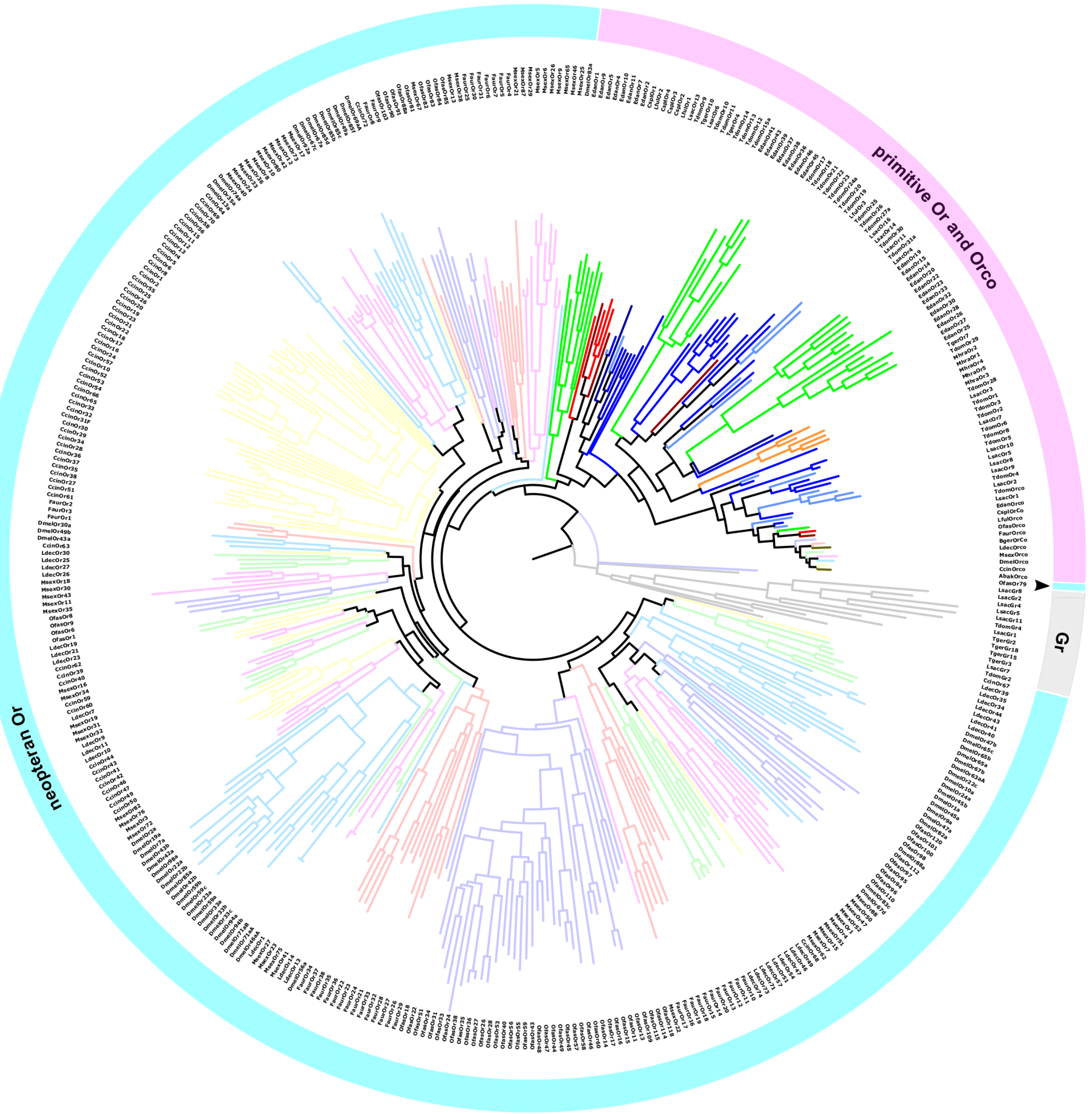


**Figure S1. Maximum Likelihood tree of insect Grs and Ors based on full length sequences.** Tree is rooted using Grs (grey) as the outgroup. Different colours represent different species. Neopteran Ors represented by desaturated colors. Species abbreviations: Lsac – *Lepisma saccharina (*light blue); Tger – *Tricholepidion gertschi* (dark blue); Tdom – *Thermobia domestica (*blue); Mhra – *Machilis hrabei* (orange); Edan – *Ephemera danica* (green); Lful – *Ladona fulva* (red); Cspl – *Calopteryx splendens* (dark red); Faur – *Forficula auricularia* (desaturated red); Ofas – *Oncopeltus fasciatus* (desaturated purple); Ccin – *Cephus cinctus* (desaturated yellow); Ldec – *Leptinotarsa decemlineata* (desaturated green); Msex – *Manduca sexta* (desaturated pink); Dmel – *Drosophila melanogaster* (desaturated blue); Bger – *Blattella germanica* (brown); Abak – *Apocrypta bakeri* (brown). Note clear separation between primitive Ors and neopteran Ors with the exception of OfasOr79 (arrowhead).

>LsacOr1
MKHIRQGLVADIWPVIKTMKFFGHYFFRYTEDDSSMKTAFRGIYSILNISLVTLHFIFGS
VSIIFKLNDIEGLVANAISTFFALHAVTKLIYFAIRRNAFYQTLDCWDVINSHPMFAESN
LRYKEIAIKKVRRLVLCVSSAILLFAAFWSSKPFFASPFREIESGNETMIVEGTLLIVDA
WYPWSLHEFTFFAASYLYQLYWLIFCMFQVNSLDMLFCAWMIYSIEQLKHLKEIMTP
LVELSAGRDPDALRKAELWPEINTIDNTVSRQDTPPSYQAAARNRIYPETLNVEMERS
MVISDFAHLKEPIIAYNSDEANIGDNVLTIKQQVYVRSAIKYWVERHKHVVRFIESVX
NTYGFALLLHMLTSTITLTLLAYEATKISAFDVYAMNVIGYLLYNLLQVFMFCIFGNNL
IEESISVMKAAYECPWYNGSEEAKTFIQIVCQQCQRALSISGAKFFTVSLDLFASVLGA
VVTYFMVLIQLK*

>LsacOr2
MKLTNLMECVWTNVRVMQYVGHFMIDYGKDQSKRKQMLHALYSFAHMSLITTHFV
FSLIDLVMKRNDLDALVSNAISTCFTVHEVTKLYYFMINRKKFRETLGMWDEKNSHP
LFSASDAKYQENAVKTIRWVXFFVLFGTEVAGSVWAIKPFIMRKTVKVQEGNATVVT
HRRMLICEAWYPVDIEDTGNFIYVYIYQLYWLAVGIFQVTSIDVLFCSWLIYACEQLK
HLKEIMVLLVELSAGRDPKALREAERQTDVNSDHHTDGRDMIISVQQAPRNRIHPTT
FNVDMYKSLDDGNVGENALTTEQQLYVRSAIKYWVERHKHIVRFIDSVGDTYGFAL
LLHMLTSTVALSLLGYELAKITALDVWAMNIVGYMIYTLAQVFLFCIFGNKLIEESKS
VLQAAYECSWEDGSNEAQIFIRVVSQQCQKPLAISGANFFTVSLELFGSVCGAVVTYC
MVLIQMK*

>LsacOr3
MKWESKGLLRILSPHITFLQFCGHFVLDFHTNKAPMLQFFRGSYSVIQISLTFLHMAF
CFLKIVFSLSDLTKLVPAVVTTMLGIHGVIKLIYVASRXKRLARTLRLWDEVKTHLIFE
RNDEETLNRTKRRAAKWLIIAAIFYAFFSVAWIISPIFDSGFEDIIVDNVTMTVEKPRLI
VGAWYPMNLTTSPGYQIAYMYQMYWAVFCPMQVYGIDILFCCLLVHASEQLKHPKQ
ILIPLVEMSSNREGNPSAKSKYLGLSQLSLFQDNSLLSELPRRRQVAWSNNKIYVDEIL
NNDNNLNAETGASPGDQGMAAQIMDAEKKKTVRSAIKYWVERHRQIIRFTVEVED
MYGVALLLHILLAAMNLCILAFESSQIRQMNIYSINVLGFMIQNLLHIFVFCIEGNNLM
EESSSLMRSVYDSSWYAGSEEAKVFIQIVSQQCQRPLSISGAKFFTLSFDFFGSVLGAVI
TYFIVLVQMK*

>LsacOr4
MITKQWCIAQVMIYPLTLAALSGHWPIRLPPNAEKIKYFTVRAIFTIVVIVVYCLMLTS
SIIDFCLNLDNITHLATNAIPTIFIAQSAFRVILLAWKRKEFSKMMLSCSLMARNCKFKI
FQNEYYLAFERQIKVMQYMDTVFRVACTLLWAVAPLLVHEVPVAHNGTQIVQEIHIM
HGWYPFDMDNPYVKDVIYAVHVTILILCTMHIMGCDYSFMTFFMFATHQQRHLKTV
LKDSLLEGLSGKYPTNVTDQLEQSKGFCAPGYIVNQISSVIDPGVAHKISQKKTDVME
MRREGILGELKEWVKCHLEVIRLVDNINQISSTLVLVTFVGNAIQLCLMPFLLIQARG
NWCIIGPIGAYFVLTITQTFLFCSIGQTVREESVSLSDALYEGNWPEASTDLNLMLLLH
MINQRLQRPLAVVGGGFFILSLDFFTSLLGAVLAYFVVLVQLK*

>LsacOr5
MDAKFKPRGLVAILWPSIRVMQLCGYLILDFYTNNSPFRVTMRVIFSVVHMAIAILQC
LSSVAAAFLNTEDFREFVFVISLSIYNINTVIKLLSISLRRMNFYKSLTLWDDMEFHPLF
KESDERSRNKTYSVNKKLLVLFNAMAVVDGLIWCAWIQIYPDMKEIEYGNSTIVVTY
RRLIIVGWYPWNIHTTIGCLISKPNRFYWMCFVIFQMTTFDITLCSCLNHICNQIRHLK
EIVAEVVELGRDQIEPKEILERNDYRESSVSEVNAVALYNANSSLDRKYLVQSAIAYW
TKRHRETIRFMGAVDDVFGFPLMLYMLLEMIAATLVVFEASKDKQMTFIANVHGYIIL

IFFGVFLFSYFGQKVIDESTSLLRTLYDTPWYDTSDEVQVLLKIVTEQCKKPLAFSGG
MFFKLSYDFFSALLGSVATYYMFLVQIK*

>LsacOr6
MAEHTAMRDMLTLNFKLLRYSGHWMIEEPRNVWFTKRKFWLTMYRSVIRVFSDCHS
ITTAIEFVLNITDIDKATRNVVIVLYNTNSFCKMSYYAFRRDTFEDIEETWNIEFKRDSN
FEEFRQSREELIEVSRKYSKALSLMITISAVSLVFKWCFFPLTVTQPPPDWKPEIIGNDT
SGAVFRVLPADAWFPFNALRSPAYEIMTFCQTVGATYIAFQGGTFDAYFIGVLIYIVAM
MKHLRWALGKLVDLSLEADENNESSSASIDEENNQLARNKNQKVDDAIVYCVQLHQ
VITRSVKEINMLFGPVVFVQFVISVLAICTVTFQATTIRGYGLKIMSLIEYLVVSLIQLF
VFCFHGNRLIIESLAITDEAYGKQWHRTPSKVKTTLRNLLLRSKKALDLRGVGMYSL
SMENFLSIVTASFSYFMVLMSFDKGDDA*

>LsacOr7
MMNREEHRSRPGLGGLFRLAIREMQFAGHFDFDYYTDYSRWKVILRYAYSSFHFGVL
FLHIILTFVDIYFKRSDFEKMIPAIATTTYGTHGVMKALVFITKKEKFQKVLHCWDNIE
THPLFVAENEKADLTTYKRSRWITALVSVLIFIDYIFWATKPLVDYPVREEKKGNETILV
KKNIMIVECWYPFDIQITGNFLAAYAYQLYWMTVCIMTVTSIDMLFCNMLIHAREQC
KYLRNTAKLIVELTLPKRNNVHEEPLLHLTPRNEQLITTDATAGVSAESEGLQAAVQR
LDSEAKEQVVSSVIEHWVNRHNSVVRYVALVEDAYGFAVLLHMLVSAIALSVLAYEA
SMGDIGLVQAGNMLGYLIHSLGQVFLFCFYGDQLMDSSAQLMRDLYGTLWYDGNH
ETRVFLKIASVQCLKPLKVTGSKFFTLSFEFLGTFIGAVVSYYLFLVQMK*

>LsacOr8
MAIKFKSRGLVAILWLNIVVFKLSGFYMMDFYPNNSLFRVTMRAILSIVNTSIVIMHCC
LYFTTAIFSTDDFNQLSILISIMILTINTVTKFMFISLRNRTFYKTLTLWDATEDHPLFKES
DERARNKTYLLNKRLLITLYVLCPLNGFLLSIWQFLQGDIKEEILTGNTTMIVAHNRL
MLEIWYPWDIDNPVGYGLTYAIQVYWLLLMIYQLVTFDITFCTMLNHIHNHIRSLKEI
VEDVVKLGSEKEELDELPGPSETNGFGHSSNNEIEEVELYNRERLSCRRFLAQSAIQF
WVKRHGEIIRFMGAVNNMYGSAVMLHMLLDTIASSLIVYKIGMDETTHFNATTHGQ
LIFIFVRTLSFCYFGQKVIDESSSLITTVYDTPWYEATEDIQTFLKVVTEHCKKPLVFSG
AMFFSLSYDFFSAFLGSVVTYYLFLVQMQ*

>LsacOr9
MGTVTVIESSSMDTEYTPRGLVAILWPNIVVLKLGGIFFMDFNANKSFLRVTMRAIFS
VAIAFLVIMQCCLYITTTVFSFDDWNKFFLIFSQAIFTIHDVTKFLFLSLRNRTFYKTVT
LWDEMEDHPLFREAGEKARHKTHSLNKGLLIIFNVLSVINAIILCFWQAVQGDIEEEI
MMGNSTIIVTHKRLLYEIWYPWDTENPIGFVLTYVLQLYWTFVMISHVATFDMTFCSV
LNHIYHQLRFLNEIVEDVVKLGSEEKNLEELPGPSETNEFDQSSENEIKEVALYNRHRL
SPRRLLAQSGIQFWVKRHVQIIRFMGAVNNLYGYAVMIHMLLDTIASTLTVFKAGKD
EEITFSPNTHGYLIFIFVRTLSFCYFGQKIINESSSLMTTVYKTQWYKATEDVEVFLRVV
TEQCKKPLVFSGAMFFNLSYDFFTAFLGSIVTYYMFLVSVQ*

>LsacOr10
MGAAVIFPALRMDTRFKPRGLAAILWPSIRIMQLCGYLILDFYTNNSLFRVTMRVIFSV
IHTSIAVLQCIFCVAAAIVRFGHFSDLVLSMAVALYTVHNVSKLLFISMRRKLFIKTLTL
WDDMEYHPLFIESDDRARRKTYSLNKKLLLIFDGMTILNAIMWAAWLWVQGDIEEKI
MAGNSTIVVRHKRLIIDAWYPWDVYSSIGFGATYALQVYWCMFIAFQVPTFDITLCS
MLNHACNQIRHLKEIVEEVVELGSGQIETEEVSGSPETNDSGNMRDNEVKVVASHRS
QTPSDSRHIVQSGIEYWIKRHYKIIRFLDTINNMYGTPMMLHMLLETIATTLTAYEASK

DEGGNFLALVHGYMITILLRVFLFCYFGQKVIDESSSLFTTVYDTAWYEATDDVHVFL
KIVTEQSKKPLKFSGAKFFTLSYDFFSALLGSVVSYYLFLVQFN*

>LsacOr11
KINNFFRKTPWTIEEPLKFLIIKAAFTGMWVPCRLKNDRINTKPLWRLYSFLFAVGAST
FCILTVVDFFLKYGNFLAMIFNCLSTLFIFEAAFKLIYIFYKRHELIRIMEVCNEMGNLA
FFPEFQSVRLKVMEHCFKITYYIFAFTCSFSAVFWVVVPFVTGDKSQWQQVIASWYPF
DYHSSPGYEISMGYQILSACTLAAVYMTTDGCFMAFFVFIHSQFSFLKMSIDKILRGSE
LRDYYNGKRDGSRLNHRRFLSLVISDVQGGISPCEELFYNVQERQNLPTMTNNNTAW
NAKERKLLEWLQVHQATIRLADDVNNVASPVVTIVFLADGLVLCLLAFMATRVKDA
VSVMSLLVYFVLILVQVYIYCLCGDKVKNESESLANAVYDSPWMDLSPENIQTLQMI
AQRCQKPVTINGGAFFTLSREFFLAVMGSVLTYFIVLMQLPDK*

>LsacOr12
SLRTFLKI*IPFKILANMFSFPTLHRCVRFNFQLLTLSGFWVSTVDDNGKRRSVLHWSF
WIAFLYRTVITLMVTVHFLAVVAGCLRNAXKKRIKGALDTWNDTYTNSTFSKSRVAA
VEASTVTSQKVSICLIVAYLTLAVQWCVAXTSVQFDXLQEVNSTFMXVTEYTRPLPFL
SWYPFDFQRSPIYELIYILQIVSAVYFALIVASFDSFFCSLMAQTVGQLDHLKSSLGFLI
DVCIDRESIPQDKTKSVTSIDEL

>LsacOr13
YRKYISMGSGRIRPDSNMYNAVVRKNNAMQTFGNREFGVLTGDYWENXKASMNYC
IHHHQYLIRFVDALEDLFAASMLVQFLYSTGLLCVLAFEATLIKGLEIKTLTLVLFLIVS
VLQLFSICGYGNKILSXSTRVTEDAYAKPWHKGSEDVKNVIQIIFQRSQRSLQLTGANI
FTVNLETFANVLAASFTYFMVLIQLG*

>LsacOr14
NTTHYRASSGINTEYMGHKVEIITSDDDSVRNEALERQSPEGSNQAGTTPLPEVLLIW
IKEHQQLTRLMDDVQEIVSPIIFATFIVSQIVLFLTAFAFIKIPDPVYASFFLSFLIYVLTQL
GILSFLGDELIQESAITPSTLYNMSFWKCNKELSTSFQIISNTAKKPFVITGLGFFKLSKE
FYSSVLGFAVSYFLVVVQLAE*

>LsacOr15
RDAKVDCIVDE*TEEEVLEALDYCVRIHQKILRYCEGLNSVFGIVVMEQFLISTLTLCT
LAFQATTIQGYGMKVLNIIEYIICATIQLFIMSRYGNHLLIESASITNEAYSRDWHQCPM
KVKGCLKLIFLRSRRRLQISG

>LsacOr16
GKSIQEYAYLEKEQHVNFSAESRVFDQLETLNIPTVAEIPKQMNLNLEQSLASWIQEH
QDLSRFVGDLEAAVSPIIFYTFFSSQCCLFMIAFATVKVDDPIYVSFFLTLGIYVVTELA
MLSILGTQLTEESFATLRHLYDIPWWSCSRGFKMAARIVNTTWNKPFALTGLGFFKLS
VDVFSSVLGVTVSYFLVLMQFQN*

>TgerOr1
GEILKSAMSYCIHHHQHMIKFVDAMEVFFAPVLLVQLLISTGLLCVVAFEATLIRGFD
MKAVNMIIFLLVSVLQLFAICYNGNRLIAESVSVTXXXXXXXXXSIFTVDLETFANVLA
AAFTYFMVLIQVG*

>TgerOr2
WSYPTLLLALVSSLFVTIHRIYLVCLYGAELIEQSQGVYLSGYNGPWYETSGNERFLL
QLIFQRAQTPLSITGAKFFTVSMELFVTILR

>TgerOr3
LSVSESAYRIPWISCHKDIKTSINFITLRSTVPARLTGCGIFSLSLDTFGSIVGAAVSYYL
VLKNFE*

>TgerOr4
VLDGLIYCVRVHAQMLRFLDELESIFSPIMLFQFVASTLVLCLLAFDASTSLQGGMRIF
NLIQYILVATMQLYLYCWFGNTLIQESLSLPVAAYRVPWFGCHNNIKTCVRLMTLRSRI
PAQLTGATIFNLSLETFKSILGAAFSYFLVLKNVQ*

>TgerOr5
INVIVERCKTPLGFTGMGYFQLSMEFYNSMLGAAVTYFLVLMSLN*

>TgerOr6
LRCWYPFNTDASPVYEMTIIFQLVASVVLVLITSSFDSFFCALLAQTVAQXXXXXXXX
XXXXXXXLEHLKGSLGFLVDVSIDRDGPVEMGKSVSSLDDLVSRTISGTGLYRRYLG
DNRNPAKPESALFNINRGGNKVQDLT

>TgerOr7
EAARSRQTAPTHTHTHGLMDEINDFTSPVVLSTFLLSVLNLCFLTYLVARNDGAVSFVI
TSVAYACNELLYMFLYCNYGENIKQQSVSLLDEVYGDPWLGAPTEITLALHTISHRCQ
RPLEVTGGNFFSLSYSFYVCVLSSILTYFIVLIQIKE*

>TgerOr8
KYAEDVKAEVLSSVVTYFIVLIQIH*

>TgerOr9
PDAEAGGNKPLEVRRLIELWILQHQHLLRLVDEIQDTFSLLIFVCSLIDVXXXXXXXX
XXXXXXXXLTAFVVARLDDPMFAVVVGGYLVTVLLQMYLYSVHGNIVTDESLLLPR
AL

>TgerOr10
FQSNAISSRRKLSEQTFASTEEAEENIRMLENHGHVQKSDLKDEELFQSFXSYIIDIHQ
RTIRFVKLVESTFAFPVLMQNTFSTTTICFIAFQAATMKEFNLKFFNNLEYLLVAMLQM
FIYARHGQRLITQSLRLTESAYRKKWHLAPPKLHGFMKVIFLNSRRALRVSGYGVFTL
SLESFASVLGAAFSYFLVLIHLQ*

>TgerOr11
LIRLWDDNDTCPDLFLKPKCDMITRARKHARHLTILILTSSLVLVTKWCFXXXXXXXX
XXXXXXXDPTRFRLLPANAWYPFDPLVTPVYQWISFYQALSADLTSAQVSTFDAWFL

>LsacGr1
MKAIDLMASFAPYYYICKPFAVFPVSRVKNEKKELSFQNSIISRIYSLVLYFIFFVPTSVL
VTYKIIQELEHYKEKRVHMLVNTFLLLVDTCPVLIASFGTFVFIVYGSRKVPSIFSRMA
HADKVLEYRLSGHVKARRLLVGKAVLVFSVLVVLFVDLMVYKLVHMKHVNLWVVL
GVNCYFALHVLNVVLDMQFVGVVSLLKGRFGHLNKEVRKLQGYKEPDRPEPRPNS
VHPDGWNERMKLPPLTAARVQNVQPPCDFRERVKSLRTIHFELCDVVEDMNVAYGF
TMLAHFCDILFSVMYCSYHFIIHVTDLFAFPSHQILFNMLEVTVVSVRTIIITWICYATS
EEANNTRNVVQKLTVETDMKSENASELRQFSSQLLHRRKVKLMACGVFQIDLALLR
SIATSVMMYLVVLIQFDIQAKHHVLIGDTPHGDSTTESSAVTTSIPIE*

>LsacGr2
MAPHKKCQVLISPVFHTTRIIGLAPLSVLKGGSSTSYQSSFLLLFYSVVYLFVLTSIYLL
RLQWIFQEVAPSNIRNVDNWTFFAKEVLLFVSGLLIGLLSLIQSFQLQTLLEKVCKVDE
TLKITESRQRKTCRFSVLYLVLYLAYFIVENVLYGLAFPYGLTWNVIAYFTMTGYWIG
VSMQFVALCREIESRFSQLSEEIFECFKDEKHCYTEAFLHELSYDVYSCEEDRAKREP
KDEVSSIEHLTNNFIQVSSCQDGVHIPKIQIESEIPDSVDTLEEIRSVYDSMCDVVDVFN
RAFGIRLLMIVLYKFFDVIFNIFCCIYKGKFMFVHHLETTLVVTAVLQVTQFFLLVFPG
NCVMAEAQKVEVVLHKLRNQRLHTCIKEEVELFSSLLKRRKVKINACGVFAIDSTLA
FWIIALVITYVVLLSQALIIGH*

>LsacGr3
RETSTSKAEKTGVLVHKLQDSGLYPAIKEEIQMFSSLLLYRKVQFTACGFFPLDLTLLH
SIAGSVTTYLVILIQFQIASKPTNDEELPESTSTEKWVSTT*

>LsacGr4
KYNFNFKYFLLPYFMSLYLGLCVLIQFMSFCCAMKERFLQLNKRIMLAEERLQDSSQ
KAFVTDCNMGEYRNVTGVLSDADTFQALRYGYDALCDTTDTLDSTYGLRMLGIVA
HSFCETVLYIFFAIHNVERLTSDALSTVALCACFLGPILINVLLTSCSSSVLSEAEKTGIL
VHKLQNRRLEPGVKEEVQMFSAQLLYRKVQFTACGFFPLDFTLLYSIAGAVTTYLVIL
IQFQVASKDQQNSTLRTDSTTESWLTSTNLTK*

>LsacGr5
VNSEKSLVKQSGYQQQFKFNNKIIPLRSTNKQEADELFLSFQNIINAYEKLCDATRML
NSVFGMCLLIYITNGFLQIVFTIFCVSRNNDFPDTAVITAVLFMSTLFPASAIILITTSSVFI
QKKAYQTGILVHKLQNRRLNPALKDKITNFCAQLLYTKITFTACGLFPLDFTLVHTIAG
AVTTYLVILLQL*

>LsacGr6
HRKIEFSICGLCPLDFTLLYYIA*AGTTYLIILVQFQTDKRHMEMSTPPS*

>LsacGr7
KLIWDAIATIILGLRCFIQFCLCQNMAKFLPTLFSALEKFGAANIAAEYRDRNRKLHPI
KRKCSIVARAIPMLITLSMLALSTFFVYDLFHEMMGGTNWNEVSLIIWFCIMNLWRTI
PLLLYMFICKHLSHDYKIIREAMKTLLRRLVANGVSVSAIKVSDESLELPEHKKDDHH
YELENLRCLHADLSSAVDVLSCSFGSTLLVEFFFSLVGTVVQLFMVISGVRLGIILYTV
TSFWLWIHVLNWTDKLSTQANGILEDLRGVPLSKLSQKCHIEIQMLMMQLIATPVNIT
AAGYFTIYKGLITSILGSIATYLIVMIQFKQQLLGLPENNATTSS*

>LsacGr8
FACAAEFKADRNDSGRCTGRKTKAMPDLFYHDLKPVFIVLQISGCFPIQHXXXXXHN
SQGEYTFAKCSLLTVLTILHYVIFVGMSGYFASSTIELFRKENREFDDLVFDAIRLIYIA
MPHIHLITFLCKSKQIARYLQKWYELQISFTKVTGIVLETNQRRNAMILVILTPVVTTL
TVALQHFIFVDLSLWHKGNTFYSITLSNYVEVMWVLICWSLTAAAKTYNEELKRSLS
APSTVPVVVLRRYRALWLQLCRLIEETGSVLCYQYATHFICIFIILTLSLYAAVSGVLDD
TTTCMKHLGFGIMALLFICQLHMMINSAHQVMQWAGNRVEDQLQDFHLRDIGERAE
NEVRGFLQTIHYRSPCLNLGSYVDINRRYVTSLFGTLVTYLVVLVQFKVGQPSTPPPP
GNTTDIEIQELMKEY*

>LsacGr9
LLGKTFLLECSHGLFVTVLHFFVIFSGISYSTILFTCINLLMIIQLAFASEEISIQANGVIR
CLRTVPTSNLNQRCQLQIHMFMMQLIGSPVKVTAAGFFTINKGLLTSILGSIATYLIVM
LQFSPQLIKLAGRNESGPIK*
>LsacGr10
ICFCIFIWTGMTGPYLVTEEILKRLGSLGVVGEGAVAALSTALHSKRFSKCLNDILAXX
XFEYAPFFLQQSSQISGMYSSIPSCIFRLFHDLQQLPQFNSGXRFQRLNELVEDLTKKTT
AISRATSLCDDKASKIRAKKRYKLTSSQSLNTSVDTIVQVSSSEDATADLSEKLSRLRK
LHADICNTVSCFNSCFNPQLLICFMAEIYSVVRSLYLAIVSFGSPVEESLSRTVYVIVHF
LGILSYVIGGQAVYQGSRLPLSSLENFPIDELDSLGRYQIQLFLTKVYKSPARYNAASV
FNIGAGLLAPIAGNVVTYLLVVLQFRPETFVNSTCEQCST*

>LsacGr11
TSFFPEDPKTFAMEFLGATGHREARMHKCEDLIVLRRCHDRLCGLIRELISVYQLQMV
MISINVXXXXXXXSRLHMEDCVMIYTYLTFVIIVFAGTTLISTMTTKEAQGSGVLLYKF
ADITLTHDVKEEIKLFSLQLLQSEPKFSVCGFFSLDLTLLHSMAAALTTYLVILIQM

>LsacGr12
NDMYFICVQLLQSLTPREGIVNAVYYFGSFACLLLRSTTVLVCGSRIYHESKKPAKILY
SCPSESYNVEVQRFLEQVLSDEVALTGMNFFPVTRKLLLTVAGTIVTYEVVLLQFNKG
QGTE*

>LsacGr13
GCLSSRTCSYTLGCKAGF*VSHKAQQIGVLVHKLQDSRLDPLIKEEVQMFSAQLMYR
KVEFTACGFFPLDFTLLHSIAGXILIQFQISSKDKPKPSPPTDSSTEEWMSHT*

>LsacGr14
KIEFSVCGLFPLDFTLLYSIAGAVTTYLIILVQFQTDERQAEMSTKPS*

>LsacGr15
SLVRWTELYQGLVPTYGQGMFDYCYVLWSFLHLGARATTLSVLASAINDEAHRPISF
VRCCSSMEYNAEVFRFEQQLNIASIGLTGCGCFVITKPFILMIVSAICTFEVILLQSSSSN
SS*

>LsacGr16
NVNRESSRLCGSCLPGMKCFHNAIDDVETQMPKSGDQCKRXXXXXXXXXXXXXXX
XNCIRAGDSSLMVSLRSEDNERLHESFRPVLVVAQLFGIMP

>LsacGr17

RILHNELCDVVDTLDDLFGFPMLAHSLEMFLWLVFVLYSIINAVMQNSILNLDWGIVN
DIIWLLMVAGRLLILSYSCHTLPGKARAVGVSISKMLRSQMEPLVKEELQLFSSQ

>LsacGr18
PSPHTAMGGTPVKLKQLSPSP*VS*IQDKTRGFLLQAQKFLEEDVGSIDTTYMSADAK
YEVQLLVQILSTKKAVINFGGFVVVNRALLTSVFSTVMTYLVVLVQF

>LsacGr19
GYSVIWFFYSLVELFFLANSSYRTTREAKRTGSLVHKLADRSLDATMKEEIHLFSMQL
LHQKLEVSVCGFFPLDFSLLYS

>LsacGr20
ISSSAWLRTYRSLYCKVHMLLQRLGGTYPYIYIFIMGFFFIVLIFACFSFVTGILQNFSPK
FWIFISAGILVITGMSAGCIYAEMVSHEAQKFLEEDIGSIDTSRMDADTKYEVLLLVQI
LSTKKAVINLGGFVVVNRALL

>LsacGr21
PKWN*NYMPVSRCNPLPAVRTSVDADEVGNLVHKLQDSRLDPSIKEEVQIFSSQLVYR
KIEFTASGFFPLDFTLLHSIAGAVTTYLVILMQFQIGSKDKSNSTQLTVSPTQKWINTT*

>LsacGr22
AIYIISGGSHQSLLRRVSHLDEDHQIPLFSMQLLHHKMKFSVCDLFPLDFTVLYSIAGA
VTTYLIILVQFENREEHIGTTASSEF*

>LsacGr23
SINKPISVRGFLQTIHYRSPCLNLGSYVDINRRYVTSLFGTLVTYLVVLVQFKVGQPST
PPPPGNTTDIEIQELMKEY*

>TgerGr1
LVANRAGALVHKIANESLDKESKEELQLFSMQLIHRKAEFSACGFFPLDFTLLYSKNGI
ASSRSRHHFP

>TgerGr2
LYCPGCALEAGTPSASTMKNVNLLWAMSPLFFISKICGTFPFSAAERGDKLIPSTCARV
YSVALFIAVTLPGSVGLFSHIAKAVQGKTQEFETETVTSIIEIASIAIASAGSFTFTVYQC
QQLPAIISRLGTVDKILKFDSTNYGDFRKHQIYQICGAIIVSIGLSGYGIWVWYDGPDII
DNLKNMFYINLLVNALVELQVVQFAVIVEDRLSALNRELFQHRESDKQLLQNTFLKA
SMVSPDETRIASGVTHVRLQTSVVSLSQLELLRQVHHEICDXGDHLNAVYGFVLLTH
MTHIFFGVFIDLYNVILCFTSHFNLDMFVHYLTSDGLWVTYYIIKLILMVQRCSAQETA
NAMGATLCKLQSRHLDPDVKEELELFSSQLSTRKIEFSACGIFSVNSQLLHSIAASVTT
YLVILLQFHSNENTTQKQETTTCGPG*

>TgerGr3
VTIIIVCFLYTQRFQHFSVLVCKILDEHIQETWSSERATEMCNIDQEEMGRNRKTNSFK
VNVTANVGSNIFPIPCYACPLSEKLDHLRKLHGAICSSVASMNRCFNPQLLVCFIVELY
MTIRGLYIVILTYCRLEEPDIHLISVFVCVVHVLLHTFCIIMLINAGEMMXXASQLPVT
YLEELPVMHLGHREIHQVQLLLQKVKSSPCSIGATGIFNIRVGLLSSIAGNVMTYLLVV
LQFELAQKSENEKTANSTIDIIMP*

>TgerGr4
RNFYASAHRTQGSACICRGGDFGFVLAEETRVIVHKLLKRSMDSETKEELQLFSSQLS
SRNVQITACGFFPLDFTLLNSIXXXXXAF*

>TgerGr5
PVSVTTARAPLKHLYQVDMLSSAASPTCQAQIQMFLARLNDSPAEISVAGFFTLNKKL
ITSMAASIATYLVILVQFRFSEQTCLPHRNSTVQSGWNETTRAWKESVT*

>TgerGr6
EEEIELLDTTSMDDKTKYEVQLLVQILSTRRANINLGGFVTINRGLLTSIFSTVVTYLV
VLIQFKVGLPTEKRQENSTSFLPSTSPGYN*

>TgerGr7
LPISVVAALSTWHLICSLNSATFRTAGGIGATTAGAVFYGNACVATVMFMRLAGRWSQ
LMLLFHEFEQSTYRWKPRKIVLKFKVITAVIMTLAAVEHVLSIMTNAYQSNSDSGX*

>TgerGr8
SSPLTEKSTAAPVKTARYEACVLVLNEMNVEDGNVNSFNDPDQGHVLYEKDRRTTSX
XXXXXXXXXXXXXXSLEVRSILQLIFQEVQQPLTISAYGYITLSVELLVSVLGIVTTYY
VVLIQLND*

>TgerGr9
KRIQVTWANDDETLSEKLDRLRRLHEKICDCVSLINSSLSPQLLIFFSLEIAMIILHAYAI
IFFLVLSSPKYTILYFTYAHLVRLIAHTVAVVSTLYKSQTTFQMSQQPMKVLLAASTSR
MKQEERE

>TgerGr10

RHLVEGDGALPSKKSMAQELDSIRRSYEELCDSLQKMNSCLNPQMLVFFTAELFMVI
THFYVIIKDGISGFQDDGYNLFFTLAKSSGRILSTLQGFPADAFGPESAIQIQLFISKLTT
CPARLSASGVFNVGTGLLAPIAGNVVTYLLVALQFNSNLAGQKNQVTTTPSDFTT*

>TgerGr11
LDLFWSFFNIGRLTALTLHPSAVVAEASRSVMIAQTLNRRAMDPMIKEQVSFFASEVS
YRNPSFSTCGLFALNNTLLYSVA

>TgerGr12
REHYSMLCSICKTVDDNISGIVLLSFGNNLYFICLQLLNGLL*

>TgerGr13
VWFFYHVIQVLAVTVVTSGTCKEANRTAVLVQKLTLLPMDLETEKQLKWFSMQLQH
RKVGFSVCGFFPLDFTLLHSIAASVTTY

>TgerGr14
LYFFHFFFQVKMFLAKFNVSPAQFSASNVFSIGTGFIASITGNVITYLLVALQFPRYNYD
SQESHGYNSIISLSKTESSTDLI*

>TgerGr15
AKELAELVWSDSNSIRFSLRDGVVPMEAK*VVKYCESCRARL*DEISKVQGSEYPHP
MKDAVSLSEKLDGLRKLHGDICTLVSSVNRCFNPQLLVCFIAQLYNVVRSLYIVIKAA
EFDNFHTNIIYQIVYKIVFVLIHFSCILIYVIAGEAVHHSSKLPLSFLENIQAENLDVVGR
YQLQLFLTKIRFSAARFSAASVFSIGTGLLAPIAGNVITYLLVVLQFQIGDCNK*

>TgerGr16
EEKREESLVANRAGALVHKIANESLDKESKEELQLFSMQLIHRKAEFSACGFFPLDFT
LLYSIAGATTTYLVVLVQMQTDTTAASPPP*

>TgerGr17
SLICGENAVVCDLCFVVKIFSLXTILDSPYIFKRTLYLLLWGSTTAYQANRTPEILSSLNS
RSLDPDSREEIKMFLMQASCRKITFTACGLFPLDFTLLVSIAGAMTTYLVILIQFQKT*

>TgerGr18
DRKST*PNVANLTLLKQFQMSPN*IYPSLIKMTDCVFSRFQKEGKSLAGPTAIDDQLGA
YLEELSLLHEELVESVESINSCFDPQLLVAFTAGLYMIVYYIYRIIAYSTSKSSEKYIFSPS
ILSDGIALVVRLLKILLIIIMGERLKTKSRCPLLAVKRARTIHLCPAAREQVQLYILKKL
MIDPVRFCASSVFSIGPGLIAPIAGNVVTYLLVVLQFQVTGRSECCEDNIPDKCNN*

>TgerGr19
NIAVCFFDVTLFLHMLVFVVQYVENFWWPILHASVYFVAQWLRMFAICFHASCLSRQ
ANRTPSVLSSIDSRSLDPDSRAEIKMFIMQTASRKIGFTACGFFPLDFTLLVSISGAMTT
YLVILLQFH*

>TgerGr20

SFPSIFTQSSFRREQTFVVHLCXQEELITSSPLFVLCLDGYRQDVYQLHRTVLENKAG
QTGVIIHKLLRRSMAPDIKEELKLFASQLLHRKIELTACGFFPLDFTLLHSIAGAVTTYL
VI

>TgerGr21
AYQANRTQAILSSLNSRSLDPDYREEVEMFLVQACSRKIKFTACGLFPLDFTLLCSIAG
AMTTYLVILIQFQ*

>TgerGr22
RTPEILSSLNSRSLDPDSREEANRTQAILSSLNSRSLDPDYREEVEMFLVQACSRKIKFT
ACGLFPLDFTLLCSIAGAMTTYLVILIQFQ*

>TgerGr23
KVQFTAYGFFPLDFTLLHTVAGAVTTYLVILIQFQSSSDREPMIQHENATVEI*

>FaurOrco
MSAKLRQPGLVADMWPLIRMMQLSGHYLFEYHTEMGGFWTGMRLGYSIIQTVLIVI
QFGFLFLNLAGQADDVNDLAANTITVLFFTHSITKFIYFAVLRQSFYRTLGAWNTINSH
PLFSESHSRYHAVAVGKIRKLILYIVIATLLSIGAWTGITFVGESTREISDPDSENETMIVE
VPRLMLRSYYPWDSMDGMGWMTSFIYQFYWLFFTLFHANLLDTMFCCWLIFTCEQ
LVHLKEIMKPLVELSAALDTLMPHPTELFRVPTGNQIQGSQNGNYEVNLRGLYSNQS
DFNGFRNTGPLAMVDHSTTIVGPNGLSRKQEILVRSAIKYWVERHKHVVRFVGMIGD
TYGGALLLHMLTSTITLTLLAYQATKIDGVDVYAASVIGYLLYTLGQVFLFCIFGNRLI
EESSSVLEAAYSVEWYDGSEEAKTFIQIVCQQCQKPMSISGAKFFTVSLDLFASVLGA
VVTYFMVLVQLN*

>FaurOr1
TSLTKCVIYHQNSLKYVEGLQDLLGPFLFFQFLSSTIILGIAIYKFAILASTPEKIAMIPY
VLNMLLQLWFHCYLGNQLTVMCDKVARFAYLTSWEIFPIKYKLTISFIIFRAQRAVQIF
GGKFYVLSIETFISILRASFSYFTLLKEMN

>FaurOr2
VLFVSFCLMYIDTIVHFNNINIVATNLSASITGAINIIKILYFLSKHKTIARVMEQLNKGL
YPGDPGPFLQKEEIRSRKRSFNLTVLMFGAAIISAISGYIKSAISLTDFDISSVPLNDTEPI
FKMMRFRQWYPKSWYHKYYNFVVFSPVIGIMTWCPTTDMGFDALIASLYIHATCQF
EIIQTRIFDYEKGNQIKDTQRYSLLKTSLIYHQSSIKLVKDLQNLLGPFLFVQFLGSTLIL
GIGIFNFAILADTTEKFVLLPYMINMLAQLWCHCYFGEQLSTQSENIANQAYLTSWEL
YPFKYKYCISLLMLRAQNPEKIYAGKFYNLSVETFRNILSASFSYYTLLKEIN*

>FaurOr3
FRSRKMEKSNLGVDRDLFGVNIKILKLVGMWNPFTRPVKKALYTIYAILVVFIIAENV
FLTYVDMFQNFSIVRVADSLPVLITISCNVIKMIYFLSRHKTLRGIINQLNLGLYPGMTI
LTEEEIRAKKRSYYLTVTVIGVGLFSCFAVYVKVGVSLYNFDMSTVSPNNTEPFMAFL
RIRQWYPTSWWFKNYVLVSLHGVVGVMLWGPPTDMGHDCLIAALLIHIACQFEVIQ
RRVADYASDEKIDDRERYELLRKCLIFHQRSLKFTERLQDLLGPFLLIQFLASTGIIGVA
IYKFAVIATTAQKLQMTSYIVNMFMQLWFHCYFGEMLTYQSSKIVNAAYYTPWENYS
NKYKFVIKYLMLKAHNPLLFFAGKFYILSVDTFRGIISASFSYYTLLQEMN*

>FaurOr4
IKQDDIDQVIRNYRECHEIGKSNILNKEMAKDLFYPWYKGSFIFNVIWLSVLIFIAIHW
FTELIMEDISNYHQNGNKDFNVSFPYVDWYFYDLKSQTGLLIAVYVSQWVTCVSNIY
FMSCFDSFIVTFMQLVCGDLQILHYQLKNLHLDHSGPNTKHTFSGIVRHHQSILNVIN
EFNTLINWPMFFNCGRIVSSVGIFIFQASQMEPGDPKMLNQFEGAAFISGQFFIYAWYA
EKLYSLNEWTAEAAYFSQWLEIEKLKKARMLSLIMQRSQKPFFFKGGMYKVNITTFL
ELLQFTFSIYTLLSQTKNKI*

>FaurOr5
IGIHWSLFPIINMVMEMISKENESTNITSKVSQNSEDNLLEKENLLLFNGWYFWEKTE
APFYYVVYFSECLVAVFMVWMVGSVDTFMYSFLLLTCAQFKQLIKSYQQINNQPNIY
FLTDNATHQLIQCMKAHQKLLKMYEEFNNAFSMVNFFHCSKTLVALCLLTFDFTTIK
AGDSKVINLVEFISIYIIEFFLYTWYTDEVTKLSGEVSVAVYECHWTEKSKPFKQSMCF
VIQRAQKRLILQGGRIFVMNLELFLGLLRLSFSIFTLLRNMNDSSLENK

>FaurOr6
PNPLAINLKVFSILGVWNLPHFPKCVIFDWIYMIFLQIMTFNVIIVTISGIIGMIIAIKEKS
DIIIPIISHFIMEILHLTNYFYLFMNKKKISKLIVSLEKCLIESENTGYGFSESFEIWKKTA
HRITKVWMFLMIYVGLHWVLAPVINYLLLAEPVHTLYHGWYPFDKNEEPAFQIIYIM
EALATMFYVYIFGSVDVFVISLLVLTAGLMEHVSNCALKMNPPIEEFGSLNIIGTKGDI
RTYKQCFRLHQKIIRFYHQLINLVSPTLFIQTCRSTCALSSLFILAVTLKTEDAKVLVYL
EYGSLYTAEFFLYTWYATEITRLGLKVADNVYLSNWQEESTSFQKSIFMMLKRSQVP
MNIPGGPFFIINLDLLIAVHRLAFSIYTVLKQLQDEPQ*

>FaurOr7
SNQTEIAYTYTLPFVGWYPYDKWSPIYYEYTYFMESVVAVAYVYYFGVTDSLIIAYLI
LCCGHLDHLLNCIRRMQERRSNESIQIYDQENVLKHQHDNNHDNEMPLEFDLTSCVK
LHREVFGMFEDFKLLISPLMLSLSLRTLCALCLFSLDFIYVKPGDAKFIILIEYLILYSLQ
FFLYTWFASEVTKMGKMVSSEIYFSNWIPASSQYRRNVSFMIMRSQKPLLLTGGPFFTI
DLGLLIQMYRTSFSIFTVLRQMKED*

>FaurOr8
MKFTDVNFDTIKGQIFLFKNLLILSFIAPSTLIAHWKIISRVVYRLLSCYVLLLSYNIIAI
AYVAFTNPDNYILMSKQLYIAIDCFAWFIFYCTLIYYRHRFFTLIEKLVEFPEPSPPFSYG
LVRKLFTYGAIYYWMNGHILLTHPIYNNIRKWLISVIYKQEEFKMVKYQSYYAWTTN
VESPLYEMHMIHYNITFLFMIIPTHSFDFIFVQIILTLSNQILMINEKLKNIEPSRNSFDEL
IELIKLHQKLLRFYNEFKLLLSPILGPLFFWRFLANIVKIFNIVLLEEFNFDFVTIIGYLFT
DTYEFFLYCWFSNHLELKLQSIEFGAYCSQWYEFDKKERILILFMMMRAEKIHPMKT
NTLFVVNMETFLWLIQEMFGNAMILRQLV*

>FaurOr9
KEVSLEYQLSKMKFKIPDLDTKAGQVWFFSLMFKLCAIWPLDKMENWHVLKKFVF
LISPPVGVIWMFVVSIGQAVALLDFPYNLFLIGEQGFVTLELMATLSLYLNLIHQKDLI
YIMLSKMDKLPEPPKKRSYKTLRSFGKGYLFIYCFNTLAFGALPLFAGGYKLPASFW
YFKQTSSPAYEIIYILEYFQGIWMIPCNVTFDFSFVHIVKFLEAQLDALADRFESVTSKS
QLIYCLKLHQDLLSIIEDFKNYLNPAFVGLFFFGCLEVCLALFEIIMMESFDATAVVVF
MIYSLSMSQFFVYCWYGSKLVKMMENISDAAYFAKWYDFEKSERLALLVLINRAQIP
MHLRASFLCVVDMKTFTAILQESLSYAMVLRQLQSAKN*

>FaurOr10
VDVFYYTILLYTQAQFKILNENLL*LXXNNNSKEEIFKNNEELNLCIKHHIQLLKISKK
LESFIGFEVMNLCLTSTCTLCFAGFNALRKFNFTIFKLSTYLFGYLTQLYIYCFLGEKLS
LKSEAVKMSAFSNNWTAKDEKYKQKLLFVMLRANFPVILKAGQFFTVSLATYSSILN
SAYTYLTILHQILN*EISYLYK*KFNAIILTSYYK**ILKK*SIFQHFSKCIRYRL

>FaurOr11
FFFSKMKRNSNFTFHTKALHLFGLWPYSNNGILYYLFTLTTLIGAIFLLVLCAYGTHLD
SELQTKLLGIIVCTSISDVFWKSCYVLWKRKNIRETIHDLQNSSLKYVKDIQDEKEKIF
NRIYKILKIRTILQVLLSNLSGLVWASKNIITDADEKTILKQFIQVPGLKKAFGFHMWF
PFAINSIPRFIFGYVIQVVLSIWVCTNFALTDLLFITMNENLKGHFDVLSYSLKNLNNPE
DPDINSKLADCLRYHDRLLGMSEKMGRLLGPQAFTQYSTFTFLFCFLGYSLLNDMGA
GLMFCIYITLLLSEVFMTSLTCDGVFQKSLEISTSAYFCCWPEFMDKSFRTGIQLCILRS
QIRPIIYRAMFSIVINMESFTRVVNSSFSYIAILQNLSE*

>FaurOr12
LNNKFPTFWNFMKFYFF*TKPLIIAYF*NI*RFHLLSSILIGSKFKSKFSSILNYLQIIFASII
SAYLALVDTFMVSLNLQLSNQFKILSYALENLTDSKDPELKQKITDCIKYHKKILKLGI
EVEKVVTLEAFSQYLMFSLLFCFLAYNISVADSLNVALYGFICYAAILLCQVFLTSKTG
DLVYSTSLSVSSSAYSCAWPEFPRPFKLGLALIIMRAQRPLTLRGLLWIPLNLETFRSVV
NTSLSYVAILRNINQ*EKLMTYLS

>FaurOr13
METKHPFQIHLKIFRILGFLAYEKNKILDIIHATFTLSGFFIWGPCIFIQSFLVEDSQLAIIS
LTLTVTFTHCGLKGLMVLRKRENIQKILEKYDYISKHFRKYYDKEMSDILNGNIVSLR
KRTIFYLCMCYMTSIPWILKGLMKDADDTVLARQKIEIPGFRRAFGAEAWYPFVIDK
YYKFIALQLHQFFTGSIILSVTYATVDCFVVALNEYLGSHFQVLGKILESIKKSSDQEM
KKKLVDCIRYHRILFELVKDVENTIGLEAFLQCFTFLFIFCFLGFTIMNANSIDQLYGIM
TYSVLFLGQVFLLTRSGNLIYENSQKIAYCAYNNEWIDSEKSFQIAILFIILRSQQTSPIK
AGYFVTLNLETFQSMVNTSFSYITILANFM*

>FaurOr14
FLVCIMVVMQLLQSLIYLLHQTKLERIVISLILMFSALTTTFRTFDMISKWNLIDDILNN
IDYLCNHFGEKLCPEEQNALHLQMKKTLKIYAIIYFYLCMCSALMWLTRPIYEILPGE
NEYDLKGHLPIENYIPWKVETRQSLILTYIFQSVSFVALCMVFGAVNPIHVSFISHVTA
QFRILNMAIDKNGIKQPWTREKLALCIQYHNQIFKLAEKVEKFSGLEIFVQCSTFVYSF
GFLGFMASLLGITNPAISRLLIYIIILLVQVYMSLKTGDNIVQESLLIGESAYFSDWHTSI
SSSQIPHIIQFIIMRSQKACMLKATSFIKVSIVTFASLINTSFGYITLLQNINKN*

>FaurOr15
MSQTKDVHKLPLKDKDRLFGFKIIHIFARLGGFRTYKKWSIFYTPFQVFIFLSLCFHAY
LQLRYSMEHFDNLEQFLIPIIILLTWSKGIMKFLIMNGYRNLLDELYDSFEICYRIYGLH
CKEEKRKLFRDLNKYMRLFSYSYILLLYITASSWIFQIFFRTLTDESRQMSKIEVADFT
WTFHVDSYFPFVIRSRSVFWMCVFFHWFNYFVMVFIVGGGDLTYVLLMKYLHGHFII
LNMEILRSRQCPNHKMKYILDKCIEYHNELFRITNKIERIVGMDILLYCLCFTILLSFSG
YLMITTHEYLNAAIMKIYISILWFIIQLYMFFKVANDIHIEMLNVSRSAYDSTWEEADLI
SKVVIKLIIMRGQRSRPMRGSRFIEMTNQTFSKLINSAFSYIAILRNVGNEE*

>FaurOr16
TINEHETPKFGLNLAYMTMHFSGLWSYESSGILYNLYQIYLPINYLLLMLLHLIAEGLF
ISSLELTVLNFIIIFTFFRMAIKFVVLIGYKSRIKKILEHFSEMYFEFALNVGEEKKKYFH
SINKTYLKSSYYYVVISVITNLLWSLDFLTKPVVEMEHQILRIEIENFTWSFPTAFWMP
FPIRSKPVFWTILAYNFMLVLWTLFIILGADFLYLQLFNYSAGHFHILGIELQKSGQLSD
KKMHESLKNSVRYQNQIYGICKDIEVICGVDIFLHCASFIIIVSFSGYLVTTIEIFSETFN
KVGVYIAFNIFQLYLIIRGGQNVYEQILNIGYNAYDSSWYNANRTNRLLILIILIRGNRS
RPLKATSFMEITKETFSRLVNSSFSYITVLQNID*

>FaurOr17
MGEKTDPNKDPKFGIKYIFLIFHICGIGKFNKPSGILYKCYQLYITLLYVTLCVLHIITAG
YYLDDLGKSTLNLIVIISHFGTLTKFTILRRNKSRIENIYTQFTRMYFDFAKNSTEEKKH
HFHLIHRSFRNFAIIFISCSIFANISWTLSFINETMDEKDRAVARIEIENFDWAFPVTFWFP
VPLRSKILFSILFLIYSQSVFTMLFVILASDFSIVLLFKYLTGHFHILGIELEKSGKLSNDE
MRRSLKNSVQYHSKIYSLCSEFEQIFGLPFLLYCMVFIAIISLSGYLVLTIDLFDSTFKK
VFSYFIFAIIQLYGVTSGGQSLFDEMNRISFCSYSSHWFDATPSNRLSILLIILRAQHSVPI
KASMFFDITKETFASLVNTAFSYVTVLRKMEMSQE*

>FaurOr18
MARKDLPEEEPRFGVPVIFRILRIGAMWPSNIYDRIYEGIMLIIIMNFALYRFIFTYNVS
QDLVELVASSVLLCTWCKGLIKYFVISRNKKRIEWIYSSFQKAYTNYGLHCKSEKYIEF
SRTNKYMDRFSYFYFCLTLCTAMSWLVQMFIRNPTEEGRALMRIEIPNFNWTFHVDT
YVPLAIHYPYEFWLWVIFHWYGNIVMVIIFAGVDPTYLLMLRYLLGHLKILNLELQRS
ASCSVHTFGEILGNCVQYHNSMFKIAKEIEVICGLDILFQCFSFILLVSFTGYLITILDLF
SPQITKIYIYIVWCNIQIYVLMKSGNDVYDEMLKVASYAYNSHWLNSSNNEKLKIMLI
ILRANHSAPLKATSFIQITKETFAKLMNSAFSYIAMLQNIDQS*

>FaurOr19
MLYIIRKNNENDPIFGLRILMKFSRFSGMSPNDNHKYLYSLFQVFLLSCFVLLFFFQIM
FTYKVLTDLEELCTSVILLITWIESLMKFSNLNWQRRRIQLIYTSFEKCYHNYGDHTG
KAKKAEFEKINREMHWFAHFYFYTTLITALTWILQMFARTLNDKSRALSAIEIPDFNW
TFHVDTYFPFVIRYQWQYLLCVLFHWNNYFWLVIVFSTVDPTYLLMLQYLHGHLRIL
GKELTLAGECAKHSLRINLFKCIDYHNELLRIAKEIEIVCGLDILLLCISFMFLVSFTGY
LITVLDLFSAPIIKIIAYLIWCNIQIFMLTKAGHDIYEEMQSIAKNAYANAWPDANKIDR
MIVMRIIQRSNQSSPMRATRFIQITRETFAGLMNSAFSYITLLR

>FaurOr20
SSELVEFLIMMRLYADVAGLLKGLYLLGVFPKENARFYYIYPIIPIMIMGSLVIQDLIYII
VSEVSLEQKSYIVVSCVSYFHAFLKHFFALFCVRKIRLIVETNEEIGELYGREYYSDFE
KRNRSIKRAKNFLFAFGCISGSTWFLGALFFPDPQIMKDQNIEIENFIGQMPLTTWMPF
KVKSKLVFYLMFTCVASSMLRTGLILGTYDTLLISLISLVAQHFMTLGNLLRKSDEKN
LVKIYKYHLRIMRLAHDVENVAQFEIFFQCCCFIFLFCFMGFTLTQKMDRLEMYDFKII
VYIFTLLFQLFIAFNAADEVQSMSSELINDVYMSEWNDPISKSRTFAVQMIMHRCQRP
LVLKSTPFITMNRENFLSIVQMGLSYIMILKNVNEMDND*

>FaurOr21
FVKRLIYGAHEKFDQQRKDEDFKEIIEKNEKNNWILCNGFAVSAQLSVLLFCIAPILKG
DNALFPNDYYFNEWYVSITKSPIYELLYFAQVVGQFLDGFLLIIGNCFIISIINYACVEFE
LLSIAFKRLSTKVEDPIDAEKILKGLIREHQELLNICDDIIYLLNPLMSVECLIISLINCTL
IFQASSVPVDNFYFFAMMIFLFCMTSELIIFCWYGTYLIEKSEEVSQSVFESHLFIKDYR
LKTFLVLLMKRAQNPVQIKAMKITIISLDLGMQLLRASYSFFTILRQLNDQE*

>FaurOr22
GWCACSLVKLRQVFSMEIEPFSFLGLLVKSIQWVGIWPFPTKTRPLYLLNLFFRLCCW
YCFVQYVLGSFANIFLTEDDISELTKIISLFLATFNQIVKAIIMVLYRDYFLEMKKNVDR
FIDNEMKDSDKNYQLIRKKYKPLNILTKCYIILLFLNGDIYCAAPLFISQPLLNGTIPIRR
LPIPMRFLWDISISPYYELTYAFEFIATPMVAVSVTVCDVFFCALIYFNSIQYECLANDL
KDIPAETNAIKILKKYVLKHRKIISFSDEIKKSIAPVVGFECVIVTIQSSFIIFAAAVMPSG
PNVFSQLIFLTTEMARLYMFGWFCSELMEKSQIGYSLYETEWYNNSKSFKTSILIMMN
KLEKPVKIVALKVILISLELCIELTRASYSFYTLISGMN*

>FaurOr23
CSSDLLFGVVMASRSSISFFGLIVHVLQWVGTWPFETRSNLLYPLVVIYRIFCMFLIVN
YFLGFFATALLSDKIEEKTDTIPLTLAVFNVMAKAAPLVIFRDYYMQFIAKVDKFIENE
MKESEKNYQLICKRSKTYNKLTKYYVIILIINAATLCIMPILGSSSTRTVNNITTVIRKM
SLPAKYPYDTNPSPIYELTFMLQLIQTPVNTVSVTISDLFYCALVYFNSIQFECLANNLK
DIPADENANRNLSKYARKHKQIIEFSLEVKKVLAPITSMECAIIVMQSSFIIFAAVLNPST

PNIVKELVFYIAEIGRLYLFGWFCSELMDKSQIGFSLYESMWYNSPKNFKTSLLIMMIK
LQKPVLITAFKIITISLELCTDLTRASYSVYTVMSGMD*

>FaurOr24
YPFDHQLPIYYELVYIFQIVSTYLTALSLTTTNLYYFALINFHSCQFECLASQMKLLLGK
QMSKDEFVKTVAHHRKILWFSEEFKNILGPIMNYQCVTVTIQSSLIIFTISTIPYGVKLF
NQLTYFFVEMGELYAFGWFCSDLLETSQIIFALYSSDWFNVPKSLKLSIWIMMMRLKP
VKIFALKIILISLELCTELTRSSYSFYTLMSRMNEKQEVE*

>FaurOr25
MMDRQLRHFHLLGLNLDLGGDTDASVVSRCFGAFLFFCCVLHNMLACGCLYSLFQE
LTLENINSLGYLIVMLLAQMDIVLVLLNRNEIRTLIGVIRDDIRQQNLIANHRGQKIAN
KYTKYLNISAKFYEYYSYLAFVVYFFVQIIRFAITGQLNLFVDIYYGNLMNNRWMYM
LIVFLNVEVSFYAMIVFPLTDMLNFALILHPACELEILSDALVNLKADAVQMLKETTGG
NLYQANLTKIMKRKLQDCIRFHQRIVKYIPKFEILYSTIFFMQFSVTFFCLCFTLFKIVL
NGISGLHAMIYLTWLLTQLYCLCACGQHLQTKSEKFYSSIFHSGWERSPIEFRAPIFIM
QERMKSPLKILVGKIFELNLDSFSVIIKGSYQIFALMMNLKE*

>FaurOr26
MPWPWEFKDGLLFTKVGLINGKERECTYSAMTLFLFTVMGIYPEVGPFWIRMLKKF
SGYMAFLSSSQLMFSLAVRLYLTRHDLELMGDSLYIFCLSTHCIYAQLLIRFNQDKISS
LFEALEDNLNAWPKRKPKEFAPPMSKNEILEKGLEYPSKTRRHPGESFLTYTLLWNNR
FMVIYTILSLGISLLYPLIPIFSEAKDGERLLPHSGFYLVNEQESPWYELLYIFLSYGASV
NAIYVPVFDCLVTVGFSKYISAHYQALRFHLESMRDASENFNFRKELIKAIGYHKNIL
KYGALVEKTFNVYLLYIYCVWVFIMAFVLLQFTMVEMYTLQFFVLMNVITVAIINLFI
ICYFPHIATEQSQALSTSLYFTDWVHTLRENKIIIYIFYNGITKPFYQTTGKYANITLDIW
TGVIKTSYSFYTLMQQMK*

>FaurOr27
MKMPWQYKNGVGYSRVLYVRGKNRLCVPTAFAVTFLTWLGLYRDSGPFWRRALKK
SIGYFAFFLSIYQFFATVSHTIIYIGEFEKMGETFYLSLICSHAIISQIFLRISGEEFGILIEKI
EDNCDAWPAREPEDFAEPKDVDEIRKEKLKAPSEQDRHPGEAIVTKTLIKNNILLFFY
WIYSCFCGTCYTFIPIIKEAKEGERLLPHDGYFIINSQISPLYEILYVIMGITAVVNALYIT
SMDMIFCNFINFITAHYQALQLRLESMHTASDNFNFRKELIKCILHHKKLKDLAEFAE
DLFNKFFLYIYLLYLLLIGFILIRFFKGTDLSTVEKVTIFNVMMISLINLFIICYFCQIITDQ
STFVPFSLYQCDWVHTNQTNRILVYIFYSGISKPIYFTTGKFASITLDIWVQLIKAGYSL
YTLFEQV*

>FaurOr28
MERSIFFSTAYLFNPKDMVLPWKFHKGLPFTKWGLLHGEKYFFLNSAFLVEAFTWLG
VYPDIGPLWMRLLKTFIGFISMIYSISCCFSSGYPLYQLEGDLYDLGDYLFLTLISFHSLL
SQIVLRTQEKRIGNLIKMMEENLFAWPVRKPEDYLPPKSIDEVRETLESPSESKPHPGE
YVLTHFILKTNWLIMFFTSSSMIVASLFGLNPFLIKNPPGVRLLPHKGFFLVDETTSPW
YELVYIWIFFGCWVLCLYGSVIDIMIYTFIRYTSVQYQAIRLRLESLLDADAQGYNIRE
QLIKCVHHHRKLSKFERYNEETFSVYYLFIYLYTLLCITFLLLKVGLTKVYNVEAFLTT
MILLILILNLYVMCYFCENVTRESLMLVNSLYQVDWINMNLETKKILIIFYYGIQKPYY
YTTGKFRKISLAMWMGVIRTGYSFFTLLFQIIE*

>FaurOr29
MASKSEKEDVIKGLTDRTSRYFRYNQILLQVIGYWPVNKDSSFWNKYSIAIQLFENIL
MVLLIISQIILVVLKWENQQLMLETIYVIMFAMAVLFKSLYLVSYAPRIVGLCVRLDER
LGVAQNWSQDQVIILEKSKNIAKRVTFALTGMFVFAGSLYVFLPLWVFWGTGGYHFQ
ENATEPDDKYFVLLNVWLPVDQRFSPGYEIGFVFKGFNAIITTFILAVGDSLVLAVIIMI
TGHFKVIQYTLENTSKFFENKNEKETINLPAQLESIMSRCIHHHQIVLEYINTCERILNI
YMMVLMSQNFLLISSALIQITLGTPSQTLPGLIGSFSCYVADLFAVCWYCNILTIETARV
SDSAWNSAWYNASSLYNKALILLMIRSKKPVRLTVAKFGPITIDTFIGVIRASYSFFAVF
NKN*

>FaurOr30
VLFRSVHAQHQRYLELPISTLRITGLLIVKYPDSNMRNLLAKIQFFFTCTVFLGAVLIQI
IGLFLGAHELKALTEVSMTMIPILMTFIKVTSTAYRVRSVRKLVDLIEDDFEQDTSTMN
QSVLNVVKHTNRFINTIFYICVFASFGYITIPVHELAQKDNTTVLDDLLPLKASYQYDL
EEERNFLITYLLETWVIINALLLVPSFDVFVVSLIAHASAQYDILKKCLENLRNEAIGNI
MMKEKNLAERPTEIEEKEDMATNIFYCNGYPIPKAKIDCEIVETLKKYINYHNRINNYI
HLLSHTFQVVFLSQIQSSLLMICFTGFQIFTQAGVNLAKTEAIIVYLTFQLLELFMFCW
SADIMSKKSEEVAQSAYSLPWSRSGSEVRFCLRMIISRAYQPVIISVAGLHTLDLQMFG
KIINTSYQFLAIMQKMAKDELYMEQMAKG*

>FaurOr31
FVSLSVTVSSDIVVVVVVECVCSLSHKMSLQFRMMRLHLQLLWLSGVPLLQQTKKR
FLNIIGKIVCFLNVETLAFLIYSQIYSIWETDDLDVTARAFIMVSAGINSIFKIPHLLLFQ
SGISKLVVQVEDTFGRKDFLLNSKYGSWIMQCVRRTNFLLILSFILAFGASSSLCLMPII
WYLNDSNLDANGSYIRELPYLCHLPFQLDCKTSPFYEFIYGHQFLMGSLAALTTIAAD
LLPITFLMHATVQLKIIGEEMQATTDKGIVIKCIKQHQKLIKFIGNVEKIFNPMLMVQF
LMSLLNFIFAAFKTINSESVGLTSNIGTLVFTIYNLWQLFIFCFFGELLMGKAISLANEG
YSSGWMEMRPLDAILVKLFIQRAQQRLVVRVAVISKLDLDTFMNILRDSYSFLAVLNT
MNN*

>FaurOr32
IFINIFTIAMLFLLALFVDIEEYKTSFINSTDLYRTDDLGMVQDIYVPWDVTEKGYFELT
WLIIMLTCSKIMTANAAANISYYYLYNEITKQLQILSYNLINIKEDALKEIDQNKIDVQ
EELDLRMRRKIKECIIHHQQIYSLTNDINELIFPSISIAWICLSLIYSFYLYLYASMEFDKD
GLLELYATLTDAMEPVPFCIIGTKLTTHSELVGFAAFSSAWYEIEPSFGRIIPLIMARSQK
YLSINTLFMGEVSLKLFSNVLQSAYSFFTVLRTVKNDE*

>FaurOr33
GNNRFHKYYRIYTYVMLSQFFITCTINLYFVFEILPVFITNLAMVFIIVDTLCKGYPIICL
REKMEPFINWMNENINRERKNESKKRQKNVENSFKKGRRVFIVYNVICGTSGFLFLA
TPALSPLLMDTPENVTIPRQLPHLCWYPFDTSPSPIFEIIYFVQIFSTYVHCMTISLMDCF
YISFVIGIISQLKGLAIGFEEINQVSSETRRNELGRLIRQHQMIIKFSESFNEFCKPMMT
MNWCAIGGILCTLIFQVSQTEIASIQFAMIMIFLTLELFHLFIFGYFGSDLVDKSFKVTH
SVYNCGWEEWQIDLKKCISFIISRAQKPLVVTAWIFGVSSLELSLALLRASYSYYA

>FaurOr34
MASFELQSYVGIQIKIMTICGFFPPNQTMRAYRFRVFYHYYMLILLAYISIFALHYAIEI
RNNINEAVRITFHGLLLLQMFIKSMSINVFYNNQFFDLIHQIDEYFIEHYPLEKHRKFIK
KHVRFIIIGCCAYNILTIVLYFLDSEHWFTGNSERPFKVPFHFPGFMELPKMLYYMVSF
SAIIFYSFFGLFCFPTVFFLQLGLFYYLIDHFQSMIVKFDTLLPENYSEKELRTSEMEKV

IIKRLVDLKKEHDKLLSAYDKLRTIITPVLIAQNICVSFLNCVLIINITSIDLAVTELAHN
VFILLICTLDHFMISWLCSKVQEMSPLLSDHVFHCGWNKMNPIFRKYLLILMSKLQKP
LIFTASVFGNVEIALTISMIQFSYSLFTILSKTN*

>FaurOr35
MIFYNKQNGYIKSLVKFMEWIGMLPLKTSDKSRYYRLRWIFYYLSNLLILLGSFSNGL
YCLLRLNEMEKEPFILYFFCVTIQQLTFFFTINYHLDLFRETIEPTIDWYFRKYANKEPH
KQWIIKHTKFIQRLSFIHFFHNLIVVIIMDLKNLSSVQRERPLPLPFDFFEKDNIPILFYY
LLLIISSFVVMSMANYEHGTLLTSLAIIIYINDHINSLKYKIERIIYKIPEKKGYFNELSVL
KHLKNFHKEHLLLMRLLDDFNKFISPWLVSINIVIGVAISLSMFSLQKGDPDTQIIMISV
SFVVSQLFIHFILSLFGSRFSSSMEEWLRELYCSGWEDLKPSSRKLILILMMKLNKPLY
LSAGIFGKMNLSLFLSVVHFTYSVFTLLSVKQY*

>FaurOr36
FSLYNPTLGYIRLQVKYFEWFGFFRPSATDSNKFYKFRLLYYFLNIVIIFLEILSVGFNN
LYFFDEPHQSPFPIYLFFLSIFQFIFFLSMNINKKTFRAVVETTVNNYFDEYVKSKQERP
RIMRNTIFIQRLFYGNVVLYVCTIYFVLSRELDTTHDERPLALPIDFFTKNYIHLYGYYI
LLTLSIMIVLWIAIFSHGVIMFNFAILVFISDHSNSMKNKIKTIIYEKSESKNEINEKSFIV
RLQSIIKQHIILQRFFKEYNKILSPWLAFLNFMIGIVYSILIMILSKPEIHMEIILVSGVVF
ANYVLIHFYWSYCGTQILYSIDGWKEKLYCSGWEDLSPKNRKLILILSMNLNKPFYLS
AGIFGKMTLDLFLSIVQFSYSLFTLLSVTQ*

>FaurOr37
MFSYKKSHGYLGLNMLYLEYIGILPPRPNTKWYKLRLVFHYFFNTAIIISAISAGLTAA
REHDNIENYSLGLYYFLAVSHEYTFVFYINYHYDILFGKIVPDVDRYFKNYSDLEPHK
KWIIKHTNLLQRFLYINRIIYFLETLVLYFKELDWQKEDRPLPLPIDLFIRYDIPLYAYYL
LLVTSILVLENMTNFYNGVFTYNFTLLIYISDHIKSLKYIIKFIIYDSEDREKMRDEKEIK
LKLMEFVKEHQNLTSFFKEYNKILAPWLGYLNIINVLLYSLLLITVYEESVGWKMLTL
AGLIFLISLSLHFIWSFYGSSYSYAMSEIKSELYSCGWENLKPSMRKLFLITMLNLNKP
LEVSAGIFGPMNLELFVLIIRFSYTVYTILSVTH*

>FaurOr38
AGAGAAGDERPMPMPVDFFLKNRLPLKGYYALFLLALLVLENMNNYYTGTVSINFS
LIIYIYDHLESLKYKIGRILYEVPGQESSFRPNWLSKYKSIVKEHNRVIRLLKEYNRIVS
PWLAFLNFTNVQFYSLIMIIIMTQDDVDMRMLSLTIAILAVLLVLHFIWSSFGSYILYSS
TELKRELYFCGWEDLDPDLRKLFLIPMMNLNKQLKLSAGIFGNMSLDLFFSLIQFSYS
LYTLLSVSV*

>FaurOr39
WPYKNSGFLYKLLQISLITTLIIQVIFQGVFTYLYFDNLTVALVALMAMLSYSKGTMKL
LFINIHNKRINHIYDSFRKAYRKFALHCQEEKKVEIDSVNKRLRRQFQIYTAICFITSVI
WSITGFVDPVTDQERFLQHIEIENFPYKFGLDFWVPFPIRSKIQFFTVYAIMVFNFFEM
AWIYGGIDCLRVSLLRYLSAHLRILGKEIEKSSECKKHSMEENLIRCVQYHKELFRITK
EIETVYRIGISVSMHYVHNSICHLRSNIE*

>FaurOr40
VCSSDLSNSQPDNMFLFKRKDVSNDGIPVFGLPENLFIAHIAGGWPYKYSGYLYKTF
QVYVFFSGIVQCFSQGVFAYVHFEELTRALVSVMLLFGYTKGLLQLFFIIINHKRIKNIY
YSFSIAYYKFAIHCQKEKEEAFTLINKRLMKQFRFYTGLCVLVVINWILVGFLEVVTEK
DRKLQGIAIEDHNFKYAQDFWLPFDISTKTQYYIVFIFLTFNMVIMISIYGGVDCLRVS

LFRYLSCHLQILAKELEKAAECDKHSMKENLIQCVHYHRELSRLAKEVETVIRYDLFL
QCCIFIVVFAISGATPNENFDSNLLKMYFVAVIHLLQIYMIMSAGADVFQDFQDISFTA
YKCPWMDKDKEIKYIIYFLIQKSILTGPLHATH

>FaurOr41
LNRCLTMEKYKPFDIHLKYFHYFGLMSSKKNGLLYNIHTITGMTLIATIGIGCMIFLYL
HGIEVINLAITISIIHAVLKGVLIIRERKFIKSIVEEFIRISNMLKYNEDKNFLILNRVETSL
KKRTVVFFYVCLITVLIWNFKNVIMNGDEETVKLQKIDIIGWKKHYAVYVWYPFVID
NQFKFYFILLYQTYSGIILAGIYAFVDSFIVALNETLGAHFEVLDNALINIKSEPEKEMR
EKLVQCIRYHQQIFSLSQRVQNFIQLEAFFHCFSFSFLFSFLGYSGIKAESMDQIYALISY
GCVIFIQLFFLTRSGDILYNNSLDIGE

>FaurOr42
SSDLENHSRYFRVNQICLRILGFWTSKHDQSRLQIILSKLFSAFVFLLMILLVISELLLCI
FNLNDTQIIFEALYVMIFALISTLKMTYLYFTVDQIISLCTLIDHHFNLNSNKWKHEQFA
ILDICLMQSKYISISIVFLFITSSLVYFCLPVYMYIFLPPIESNLTGQTDFLLFNIWTPWNQ
SETGYFEISLAFKLLNCVGTTGSAGTCDALILSIVIHVTGHFKVIQWSGKNIKEIAMMN
LNIEHEIENENEHFAIPNEMHNVFKDMIIHHQTLMRSIIKMEKLFNVYSLLQIGNIFIFIC
LILLESIIFI

>FaurOr43
PVFGLPENLFIVHIAGGWPYKNSGYLYKMYQIYIISTMTIQILCQGVFTYLYFDHLSIAL
VAFMAMLSFLTGTMKLLFINIHYKRINHIYDSFRNAYRKFALHCQEEKKVEFDSVNKR
LRGQFQIYMTICFIAALTWNISGFVDQVTEQDHILQRIEIDNFHFKFGVDFWLPFVIRS
KLQFYIVYAIFVINTFENSSIYGGIDCIRVSLLR

>FaurOr44
NLFIAHIAGGWPYKNSGYLYKTFQIYILISMTLHTISEGVFTYLNFDDLSVSLIGLMLV
LSFSKGTIKLLFINIHFKRITHIYDSFRKAYLKFALHCQEEKKMEFNSVNKRLRSQFRIF
MAICFITSMIWSITGFIDPVVEKDHILQRIEIENFSFKYGINFWLPFVIRSRFQFYIVGFLL
NINLYELTWVFCGIDCIQVSLLRYLS

>FaurOr45
VLPVIITITCNSTKMIYFLCKHKTVKRLIGKLNQGLYPGNLKFSEKEIQAKKRSSNLTAI
IVGSAIFSGFASYLKVGFALYNFDLSSVPLNDTKPFMALLRIRQWYPRSWLFNHYQIIS
LHGTVGVMTWGPVTDMGNDCLIATFLIHIACQFEMIQERIADYGTKMQLDDRKRFD
LLRKCVVFHQNSLKFTTELEQLLSPFL

>FaurOr46
QFSTSFFGLCLTLFMILQDGFTSDIHTISYLIWMYLQIFAYCILGERLISKSDNLQEAIFH
CGWERDPIRFRMIILMMLNHNKKPLQMTMLKLLTLSFETFSFILKTSYQVFTLLMNY
KSIEEE*

>FaurOr47
ETFTIITFLIVCFSIFIFELFVFCWFMEDLSIQSSAILSSVYESFWYEGSNELKKSMIIFMI
KAQKPVILSMGKFGPITMETFTTVVRGSYSFFAVFSSL*

>FaurOr48
*LMSPFKGIANLMSNRFIGNVEKIFNPMLMVQFLMSLLNFIFAAFKTINSESVGLTSNI
GTLVFTIYNLWQLFIFCFFGELLMGKAISLANEGYSSGWLGMWALAV
